# Dynamic Network Curvature Analysis of RNA-Seq Data in Sarcoma

**DOI:** 10.1101/2022.03.09.483487

**Authors:** Rena Elkin, Jung Hun Oh, Filemon Dela Cruz, Joseph O. Deasy, Andrew L. Kung, Allen R. Tannenbaum

## Abstract

In this work, we utilized network features of cancer gene interactomes to cluster pediatric sarcoma tumors and identify candidate therapeutic targets in an unsupervised manner. RNA-Seq data were mapped to protein-level interactomes to construct weighted networks for mathematical analysis. We employed a geometric approach centered on a discrete notion of curvature, which provides a measure of the functional association between genes in the context of their connectivity. Specifically, we adopted a recently proposed dynamic extension of graph curvature to extract features of the non-Euclidean, multiscale structure of genomic networks. We propose a hierarchical clustering approach to reveal preferential gene clustering according to their geometric cooperation which captured the characteristic *EWSR1-FLI1* fusion in Ewing sarcoma. We also performed *in silico* edge perturbations to assess systemic response to simulated interventions quantified by changes in curvature. These results demonstrate that geometric network-based features can be useful for identifying non-trivial gene associations in an agnostic manner.

## 1 Introduction

Pediatric sarcomas (PSs) are a diverse group of childhood cancers that are typically diagnosed based on immunohistologic features and clinical history [6]. When the clinical and histologic workup do not unequivocally determine a diagnosis, further time-intensive molecular characterization is needed to ascertain the correct classification [28]. The delay in a definitive diagnosis hinders time-sensitive decisions toward treatment planning and management. Therefore, there is a significant need to develop novel methodologies to accelerate the timeline for identifying PS subtypes. Moreover, although the genetic drivers for some PS subtypes have been described [3], oncogenic driver mutations, like the canonical *EWSR1/FLI1* fusion gene characteristic of Ewing sarcoma, have not been amenable to direct targeting and are therefore undruggable [15, 21, 22, 24, 29, 33, 34]. Thus, understanding the pathways required to maintain the cancer system is also pivotal to the identification of existing drugs that can indirectly target the drivers of these tumors.

The goal of this study was two-fold: to distinguish PS subtypes from tumor tissue RNA-Seq gene expression profiles (GEPs) and identify actionable candidate targets for therapeutic intervention. To this end, the focus of the work described in this paper is to design a classifier for identifying PS subtypes and to develop a framework for investigating the functional relationships between genes or their products. Machine learning techniques such as agglomerative hierarchical clustering methods [13, 35] and random forest models [17, 35] have had success in classifying sarcoma tumors and statistical analyses of differential patterns in gene expression (or methylation) between subtypes have been particularly useful for identifying novel biomarkers. In this work, we exploit functional network properties that consider the topology (connectivity) of biological networks in conjunction with gene expression to address each objective. More specifically, we employ a geometric approach, namely ***curvature***, rooted in the theory of optimal mass transport (OMT) [30, 31], which has not been fully explored in the context of weighted cancer networks. Curvature defined on a graph in this manner is related to the feedback connectivity, i.e., the number of invariant triangles [5]. Such a geometric, functional network representation allows for novel insight that is not apparent from genomic data alone.

Gene regulatory networks are naturally represented as weighted graphs where each gene is represented as a node (vertex), with edges between nodes representing direct interactions at the protein level, and the strength of the interactions is characterized by the corresponding edge weights. In addition to direct connections, understanding how to account for indirect cooperation is essential. For instance, it is likely that a set of genes (proteins) that form a connected subnetwork function together and are involved in a specific phenotype [14]. However, identifying relevant subnetworks in complex biological networks remains challenging compared to identifying individual key biomarkers. We will consider a method for treating this key issue that utilizes a curvature-based approach founded on notions from OMT theory in combination with network dynamics.

One can consider the weighted network as a Markov chain and look at certain graph theoretical properties such as random walks. Of particular interest is the notion of Ricci curvature between two nodes on a graph. In a continuous setting, curvature is a measure of how the local geometry deviates from Euclidean space. Intuitively, curvature is characterized by the degree to which geodesics (local paths of minimal length), obtained via parallel transport, will tend to converge or diverge in the space [7]. A standard example of a positively curved space is a sphere whose geodesics trace out the great circles. To see that the sphere has positive curvature, consider a point *p*, two tangent vectors *v_p_* and *v_pp′_* at *p* perpendicular to each other and another point *p′* obtained by parallel transport of *v_p_* in the direction of *v_pp′_* which results in the tangent vector *v_p′_* at *p′*. Then the geodesic from *p* in the direction of *v_p_* will converge with the geodesic from *p′* in the direction *v_p′_* at the top of the sphere (Figure 1). In the context of networks, curvature reflects the connectivity and interdependence between nodes. Several notions of discrete Ricci curvature applicable to graphs have been put forth [4, 10]. In this work, we use Ollivier’s formulation [23], which we simply refer to as *Ollivier-Ricci curvature,* due to its connection with OMT and favorable theoretical properties [27]. On a graph, curvature provides a measure of network robustness/fragility where positive curvature indicates positive feedback (i.e., triangles) typical of hubs while negative curvature indicates bottlenecks in communication or transfer of information typical of tree-like topologies.

**Figure 1:**
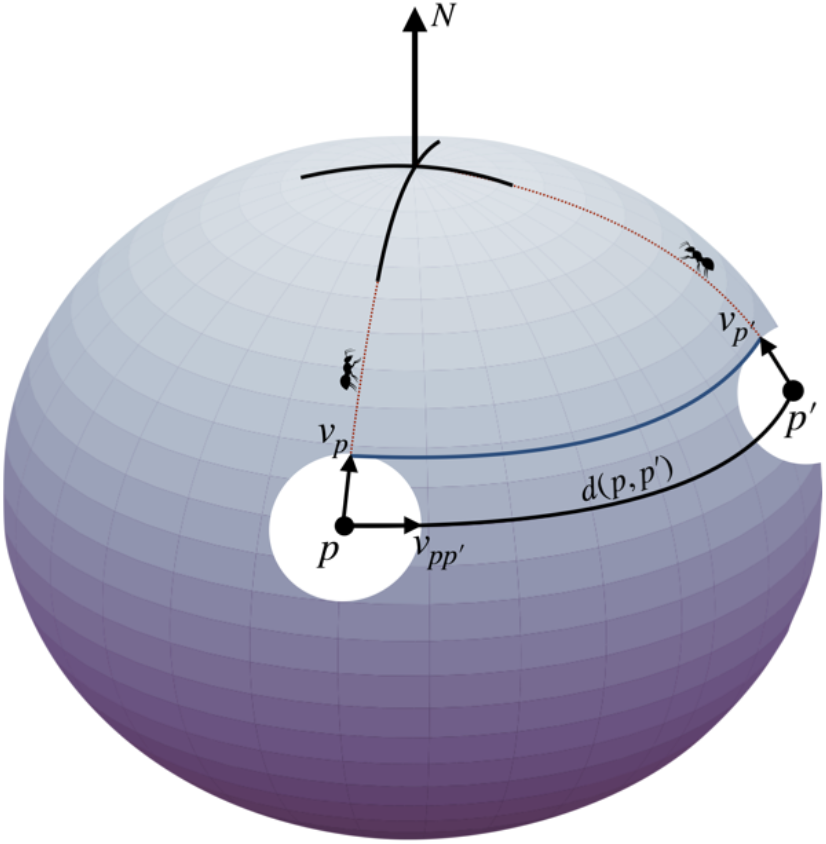
Geodesics obtained by parallel transport converge on a space with positive curvature.

In the present work, we utilize a recently developed dynamic formulation of Ollivier-Ricci curvature [11] which measures the curvature as a function of time while information is diffused throughout the network to explore the multi-scale structure of genomic networks and, specifically, to identify key subgraphs as well as the bridges connecting them. “Time” in this context is a purely numerical construct used to connote the gradations of the network organization and is used interchangeably with *scale.* The motivation is that networks exhibit varying levels of organization at different scales. Thus, persisting communities (with many connections among genes) and emerging bridges (with few connections) may identify mechanisms of resistance and actionable targets for intervention. This dynamic notion of curvature is particularly attractive for gene regulatory networks that typically have strong hub nodes and low modularity, which is challenging to overcome with standard community detection approaches. Moreover, Ollivier’s notion of Ricci curvature is relevant to studying network functional stability because an increase in Ollivier-Ricci curvature (resulting from an external impact exhibited by a change in interaction (strength) between network components) is positively correlated to an increase in system robustness [19, 32]. The connection between curvature and network robustness/fragility is linked by entropy [19]. However, unlike entropy which is a nodal attribute and thereby exhibits a loss of information by construction due to a weighted contraction of edge dependencies, Ricci curvature is an edge attribute that preserves such geometric quantities. The significance of this theoretical result has been demonstrated on real networks supporting the use of curvature as an indicator of network robustness [9, 27]. Thus, curvature concurrently computed with *in silico* experiments simulating gene knockout or pathway interference is performed to assess the network response to targeting the key contributors to gene signaling dysregulation in the cancer network identified by the multi-scale dynamical analysis, with particular attention in this work given to Ewing sarcoma (EWS).

## 2 Methods

### 2.1 Data

RNA-Seq data were generated from tumor tissues in PS patients who were treated at our institute. RNA-Seq data were preprocessed using rlog normalization. This study was approved by Internal Review Board at Memorial Sloan Kettering Cancer Center. The patients provided their written informed consent to participate in this study. In total, the cohort consisted of 102 samples from 21 different subtypes that were predominantly sequenced from metastatic or relapsed tumors. In this work, we considered the 70 samples from the four largest subtypes: osteosarcoma (OST; *n* = 29), desmoplastic small round cell tumor (DSRCT; *n* = 20), EWS (*n* = 12) and embryonal rhabdomyosarcoma (embryonal RMS; *n* = 9) and concentrated on the EWS cohort for functional analysis. The criterion for gene inclusion was a minimum of 10 samples with 10 read counts in RNA-Seq data.

### 2.2 Graph topology

The network topology was derived from the Human Protein Reference Database (HPRD) [25, 16]. The graph was then constructed by restricting the set of genes in the given dataset to the HPRD and extracting the largest connected component network, resulting in a simple graph with 8,127 nodes, 32,750 edges, and an average degree of 8.1 after removing multi-edges and self-edges. 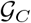 is used to denote the graph used for tumor clustering while 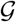 is used when referring to an arbitrary graph, which is assumed to be simple, undirected and connected. As the nodes of the graph refer to genes, the terms *gene* and *node* are used interchangeably.

### 2.3 Unsupervised clustering

We follow the construction of forming a Markov chain based on RNA-Seq gene expression data from [26]. The Ollivier-Ricci curvature is defined on such a Markov chain as we will describe below.

Gene expression data *x* ∈ ℝ^*n*^ for each sample is mapped to the graph 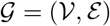 where 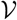 denotes the set of *n* nodes and 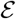 denotes the set of edges by assigning node weights *w_i_* = *x_i_* for all nodes 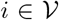. Treating the weighted graph as a Markov chain, the probability of going from node *i* to node *j* on a random walk is expressed as

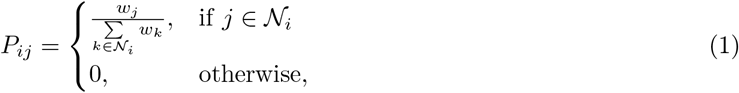

where 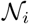 denotes the neighborhood of node *i*: 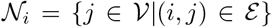. The random walk on 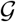 with a transition probability matrix *P* corresponds to an irreducible Markov chain since 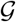 is connected. This along with the Perron-Frobenius theorem for nonnegative matrices guarantees the existence of a unique *stationary distribution π*, which is the probability distribution defined on 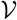 that satisfies

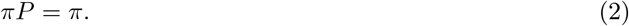

The stationary distribution may be efficiently computed from its closed form

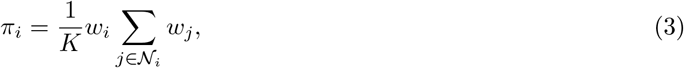

where *K* is a normalization factor.

The stationary distribution is the limiting behavior of a random walk on 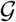 and the value *π_i_* of its *i*-th component is related to the relative amount of time a random walker spends at the corresponding node *i*. We expect that the stationary distribution encodes subtype specific relative node importance and therefore expect that stationary distributions associated with transition matrices constructed from gene expression data of samples with the same subtype would be more similar than those associated with different subtypes. This motivates the use of the *Wasserstein distance W*_1_, the metric associated with OMT which gives a rigorous notion of the “shortest distance” between probability distributions, to compute the distance between stationary distributions as a measure of similarity between the corresponding samples. The Wasserstein distance between two discrete probability distributions *μ* and *v* on ℝ^*n*^ is formally expressed as

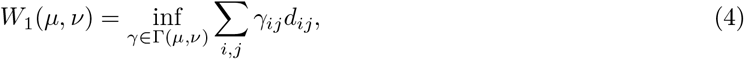

where Γ(*μ, v*) denotes the set of joint probabilities on ℝ^*n*^ x ℝ^*n*^ with marginals *μ* and *v* and *d_ij_* is the prescribed distance between the corresponding genes *i* and *j*. For details on the Wasserstein distance, more general formulations and its connection to OMT, see [30, 31, 1].

The unsupervised Wasserstein-based clustering of the samples proceeds in the following manner: invariant distributions *π*^(*s*)^ are computed for each sample *s, s* = 1,…,*S* where *S* is the number of samples. The sample-pairwise Wasserstein distance matrix **W** ∈ ℝ^*S*×*S*^ is then computed where **W**_*qr*_ = *W*_1_(*π*^(*q*)^, *π*^(*r*)^) is the Wasserstein distance between the stationary distributions associated with samples *q* and *r* using the hop distance as the graph metric dj. Hierarchical clustering is then performed using **W** as the distance matrix.

### 2.4 Geometric network analysis

#### 2.4.1 Graph construction

The graph for functional analysis 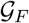 was constructed by extracting the largest connected component from 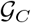 restricted to the set of genes provided by the OncoKB database [8], resulting in a simple graph with 675 nodes, 2,667 edges and an average degree of 7.9. Note that the analysis performed in this work may also be applied to the full 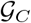 as well. The constricted network of established oncogenes and tumor suppressor genes was opted for to reduce the computational burden.

For each subtype, the strength of interaction on an edge 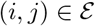, denoted 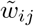, was computed as

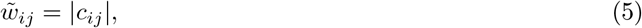

where *c_ij_* is the Pearson correlation between the corresponding genes *i* and *j*. Pearson correlation is known to be sensitive to outliers so a de-sensitized correlation was computed where samples that drastically affected the correlation value were removed. Mapping the interaction strengths 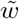 to edge weights, as described next in Equation 8, on the fixed 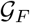 topology yields the subtype-specific weighted graph.

#### 2.4.2 Graph distance

Unless specified otherwise, the graph distance *d* is hereon assumed to be the *weighted hop distance d^w^* (i.e., *d* ≡ *d^w^*). More specifically, denote by *p^ij^* a path between nodes *i* and 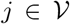 by the set of *m* + 1 nodes connecting them, i.e., *p^ij^*:= *i* = *v*_0_ ~ *v*_1_ ~ ⋯ ~ *v_m_* = *j*, where consecutive nodes *v_k_, v*_*k*+1_ ∈ (*k* = 0,1,…, *m* – 1) correspond to an edge 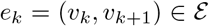 and each node only appears once. Denoting the set of all possible paths between *i* and *j* by 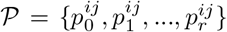 (this set is finite since the graph is finite), let 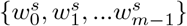 be the set of edge weights associated with path 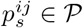 where 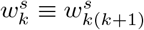 is the weight for edge 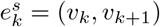. The corresponding length of the path is then expressed as

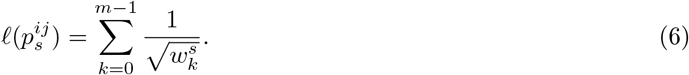

The weighted hop distance 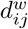 between nodes *i* and 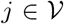 is the minimal accumulated edge weight among all paths connecting *i* and *j* formally defined as

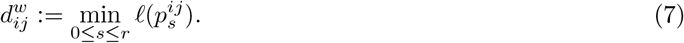

The graph under consideration is assumed to be simple, connected and undirected so at least one path is guaranteed to exist between any two nodes *i*, 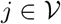. For each edge 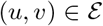, the edge weight *w_uv_* is taken to be

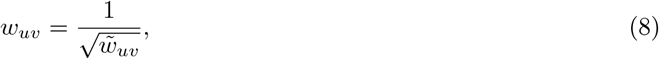

where 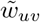 was previously prescribed in Equation (5).

#### 2.4.3 Ollivier-Ricci graph curvature

Treating a graph as a metric measure space equipped with a graph metric *d* and probability measures *μ_k_* at each node 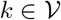, Ollivier’s [23, 32] coarse definition of curvature between any two nodes *i*, 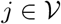 is expressed as

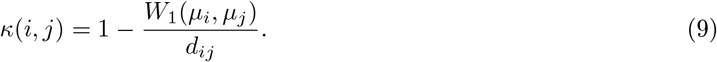

One possibility is to take the distribution *μ_k_* to be the probability of a 1-step random walk staring at node *k* given by *P_k_*, the *k*-th row of the transition matrix *P* in Equation 1. Alternatively, distributions based on lazy walks or edge weights may be used. As mentioned previously, Ricci curvature on a Riemannian manifold can be assessed by the local tendency of geodesics to converge (positive curvature) or diverge (negative curvature) [7]. Put another way, curvature may be characterized by the ratio of the distance between geodesic balls to the distance between their centers: positive (respectively, negative) curvature is characterized by the distance between geodesic balls (on average) being closer (respectively, farther) than their centers. The ratio is balanced, meaning the distance between geodesic balls is the same as the distance between their centers, in *flat* space, e.g., Euclidean space. In Equation 9, Ollivier replaces geodesic balls centered at a point with distributions supported on a node’s neighborhood and the Wasserstein distance is kindred to the distance between geodesic balls. Thus, analogous to Ricci curvature, Ollivier-Ricci curvature is characterized by the ratio of the distance between neighborhoods to the graph distance between the nodes the distributions are centered on.

#### 2.4.4 Dynamic curvature

In this paper, we employ a multi-scale extension of the Olliver-Ricci curvature on weighted graphs [11] to identify robust and fragile components of the genomic network that are obscured by the complexity (nonlinear, non-Euclidean) of the network representation. The multi-scale functional organization is captured by replacing the random walk *μ_i_* with a network diffusion process *η_i_*(*τ*) as a function of scale *τ* ∈ [0, *T*] seeded at individual nodes *i*, expressed as

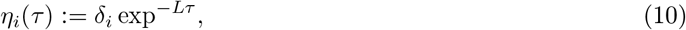

where *δ_i_* is the Dirac measure at *i* such that *δ_i_*(*j*) = 1 for *i* = *j* and 0 otherwise, and *L* = *I* – K^-1^*A* is the normalized graph Laplacian. To construct *L, I* is the *n* × *n* identity matrix where *n* is the number of nodes in the network, *K* is the diagonal degree matrix where *K_ii_* = ∑_*j*_ *A_ij_* and *A* is the weighted adjacency matrix. In this work, *A* is defined as

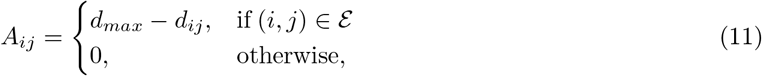

where *d_max_* = max_*ij*_ *d_ij_* is the largest distance. Accordingly, Gosztolai and Arnaudon [11] define a *dynamic* version of Ollivier-Ricci curvature as

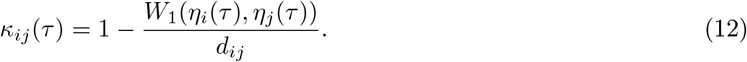

Notice that initially, the dynamic curvature is 0 at *τ* = 0 (*κ_ij_* (0) = 0) when no information has been shared and the nodes are independent. Then when the measures diffuse to steady state *π*, and the diffusion processes have completely mixed, one gets that *κ_ij_*(*τ*) = 1. The key idea, as the authors argue, is that the characteristic scales should be related to the overlap of pairs of diffused measures (a.k.a. the mixing rate) over the network. This is used as a measure of information propagation on the various subnetworks. Indeed, they derive an upper bound on the mixing time of the diffusion pair. Thus, information shared to “communal” neighbors is reflected by clusters with positive curvature at early times, whereas negative curvature is characteristic of inter-community connections (bridges) with restricted information exchange (Figure 2).

**Figure 2:**
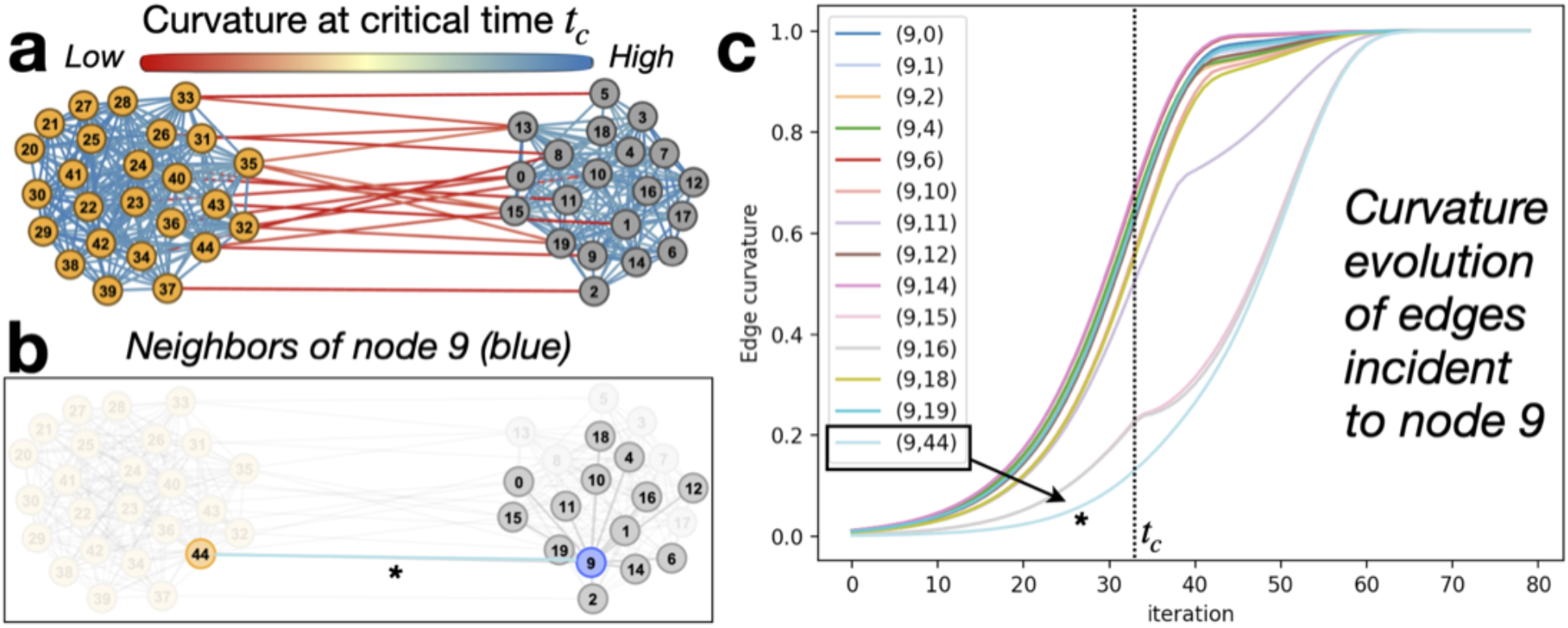
Utility of the dynamic curvature framework illustrated on an idealized stochastic block model network with two communities. (a) Bridges between clusters characteristically have negative curvature (red) while edges within clusters are positive (blue). (b) Multi-scale functional organization exhibited for node 9 is encoded by (c) the curvature evolutions of incident edges, seen by the largest gap obtained in the evolution of the bridge edge (9,44), denoted with an asterisk, that connects the two communities.

#### 2.4.5 Critical curvature filter

In addition to the multi-scale representation, there is also a hierarchical aspect within a fixed scale, as curvature measures the strength of the functional connections. The first scale that the dynamic curvature of an edge reaches a critical value, here set to 0.75, is called the *critical scale t_c_*, i.e., *κ_ij_* (*t_c_*) = 0.75. The critical scale based on this critical value is not arbitrary; it is related to the scale at which information has sufficiently diffused throughout communities but has not crossed bridge (bottleneck) edges, and is therefore an ideal scale to capture functional subnetworks [11]. Bridges may be identified as edges with negative curvature at the critical scale. Connected components that emerge by removing these bridges characterize communal affiliation amongst the nodes. Moreover, iteratively pruning edges by the critical curvature value in increasing order reveals a hierarchical structure of the functional association between nodes.

#### 2.4.6 Multi-scale functional clustering

Incorporating information from multiple scales in the dynamic range lends additional information for characterizing the intricate fabric of the network and its key sub-structures. In order to utilize the mult-scale information, we define the *average critical curvature* 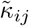 of an edge (*i,j*) as the average curvature over the critical dynamic range, expressed as

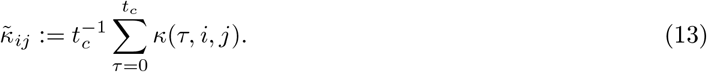

In this manner, 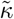 provides an enhanced measure of the interaction between nodes. The edges of the network are then iteratively pruned by their 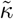 value, starting by removing all edges with negative 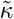 and then proceeding in increasing order. We keep track of the number of iterations nodes *i* and *j* are found in the same connected component, denoted *R_ij_* for every two nodes *i*, 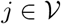 in a *persistent components score matrix R* ∈ ℝ^*n*×*n*^, where *n* is the number of nodes. With the rationale that the longer two genes remain in the same connected component, the stronger their functional association, and the “closer” they are to each other. Accordingly, we construct a gene-pairwise distance matrix between nodes *D* ∈ ℝ^*n*×*n*^ where

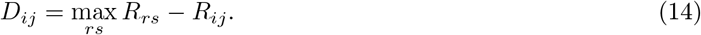

Hierarchical clustering of the genes is then performed using *D* (Equation 14) as the distance matrix. This process of hierarchical clustering based on how often nodes are found in the same connected component while iteratively filtering out edges by the average critical curvature is illustrated in Figure 3 and is referred to as hierarchical-acc.

**Figure 3:**
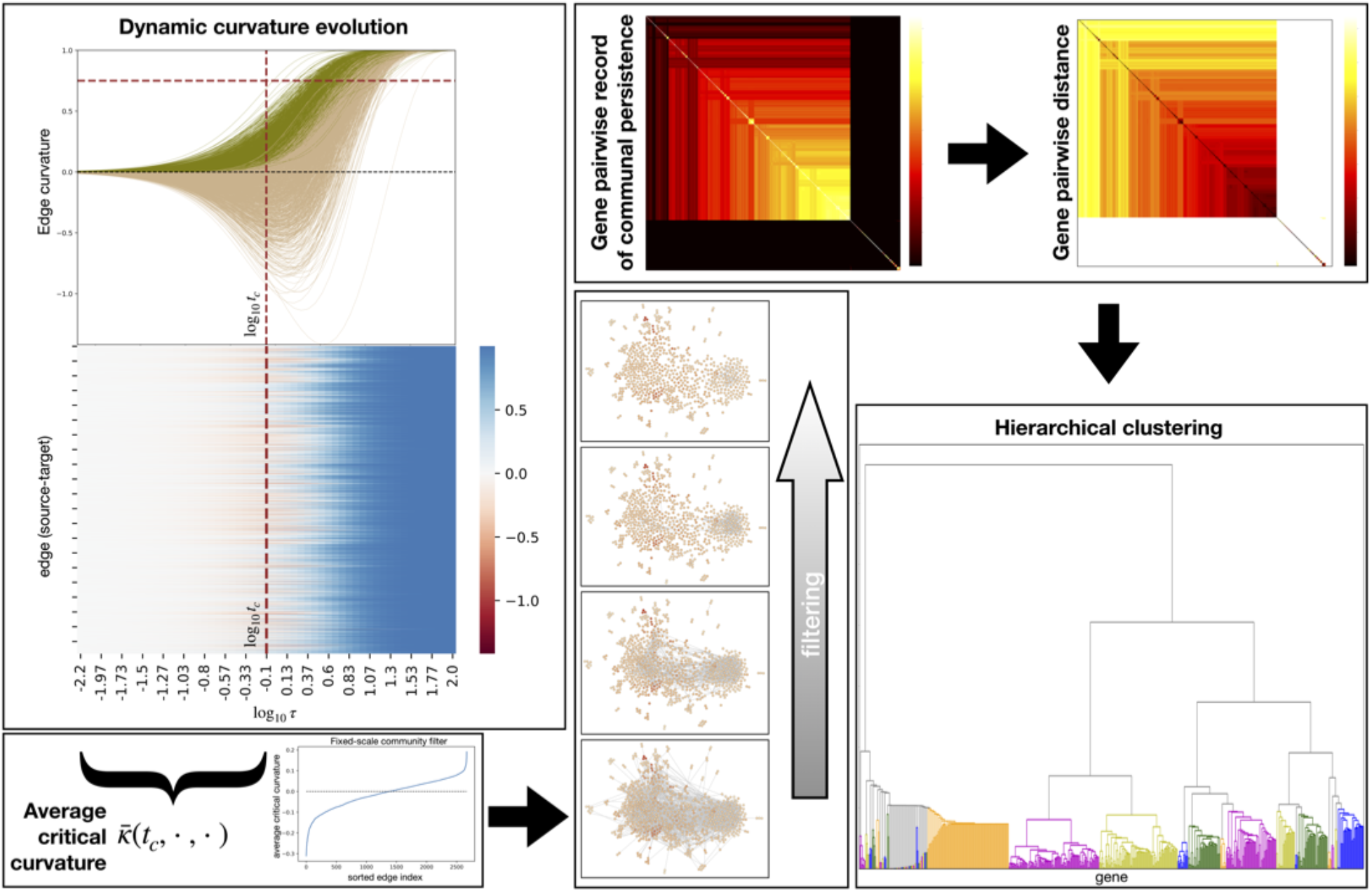
Hierarchical-acc dynamic curvature clustering pipeline.

#### 2.4.7 Perturbation simulations

To assess the network response to targeting a particular edge, curvature is re-computed while dampening a specific edge-weight. Specifically, for a fixed edge 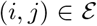 with interaction strength 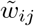 (Equation 5) and weighted hop distance *d_ij_* computed from edge weights *w_uv_* according to Equation 8, the baseline curvature between any two nodes is computed according to Equation 9. The nodal measure *μ_r_* used for the baseline curvature computation is expressed as

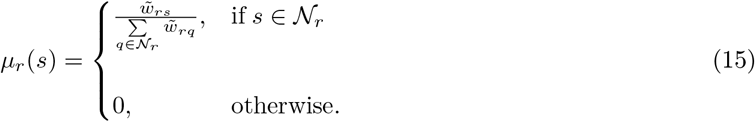

The edge perturbation procedure then proceeds as follows. The interaction strength 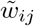 is perturbed toward 0 to simulate a disruption in communication, or cooperation, between the nodes. Our interest is to see the trend in curvature due to the simulated reduction in cooperation. To reduce the computational time, we therefore choose a coarse discretization of the interval 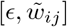 (where *ϵ* is a negligible amount, 1 x 10^-6^) into *N* = 6 uniformly spaced points: 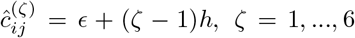, where *h* is the discretization step 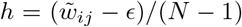. For each *ζ* =1, …,6, the perturbed edge weight 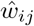 is computed as 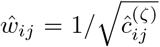 and the weighted hop distance is recomputed. Accordingly, we consider the “perturbed” probability measures 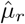 attached to node 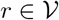 expressed as

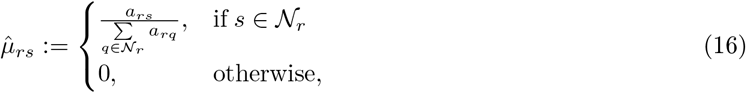

where the edge attribute *a* is defined as

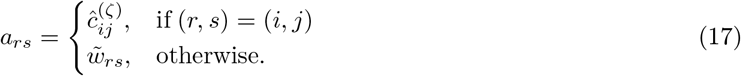

Finally, the *perturbed* Ollivier-Ricci curvature is then computed between any two nodes according to Equation 9.

## 3 Results

### 3.1 Clustering

Wasserstein based unsupervised hierarchical-acc clustering was applied to cluster samples from four PS subtypes using the whole HPRD-derived graph 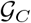. The resulting clustering was highly consistent with the histological subtypes and is shown in Figure 4 with the heatmap of the pairwise Wasserstein distances. Discarding the single embryonal RMS sample outlier which did not cluster with any subtype, the remaining 69 samples were separated into four clusters with only one misclassified sample for the histological subtypes, yielding a classification accuracy of 0.99. Of note is the incorrectly clustered EWS sample (green). While the outlier EWS sample was molecularly confirmed as a Ewing sarcoma, the case presented as an intra-abdominal soft tissue mass which is clinically atypical for this sarcoma subtype. The patient received conventional chemotherapy consistent with the Ewing sarcoma diagnosis, but demonstrated treatment-refractory disease. Considering that the methodology is agnostic to the histology and clinical classification, this serves as compelling evidence that the proposed approach will be helpful to understand why this tumor did not respond to upfront chemotherapy, possibly because its underlying biology diverges from EWS.

**Figure 4:**
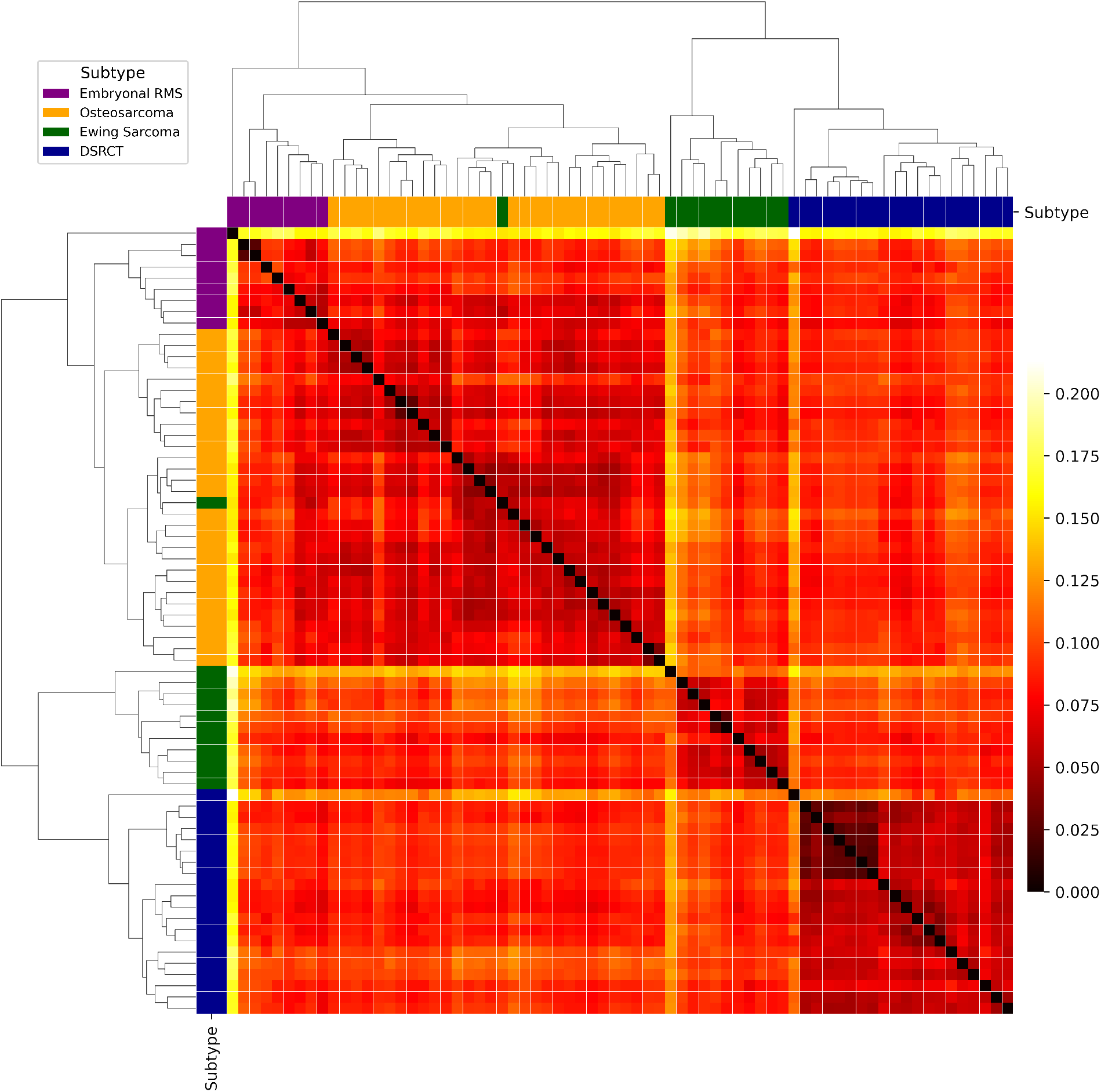
Hierarchical-acc unsupervised OMT-Wasserstein based clustering of samples in four PS subtypes using network properties on the whole HPRD-derived graph 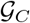.

### 3.2 Functional analysis

#### 3.2.1 Critical curvature filter: results

Preferential community formation by filtering edges with negative critical curvature (at the determined critical mixing scale) captured the characteristic *EWSR1-FLI1* fusion and the novel *FLI1-ETV6* interaction. Finding this persistent *EWSR1-FLI1-ETV6* relationship was purely a mathematical discovery with great biological significance. This was distinctly different from the connectivity between these genes found in the OST and DSRCT networks, highlighted in Figure 5. Removal of edges with negative critical curvature resulted in 1,481 (55.53%) remaining edges in the EWS network, 1,483 (55.61%) edges in the OST network, and 1,479 (55.46%) edges in the DSRCT network. By incrementally filtering edges by critical curvature value, we found that *EWSR1, FLI1* and *ETV6* form a single connected component that persists until only 587 (22.01%) edges in the network remain before *ETV6* breaks away. The *EWSR1-FLI1* association persists further until 296 (11.10%) edges remain and a majority of the network has been decomposed.

**Figure 5:**
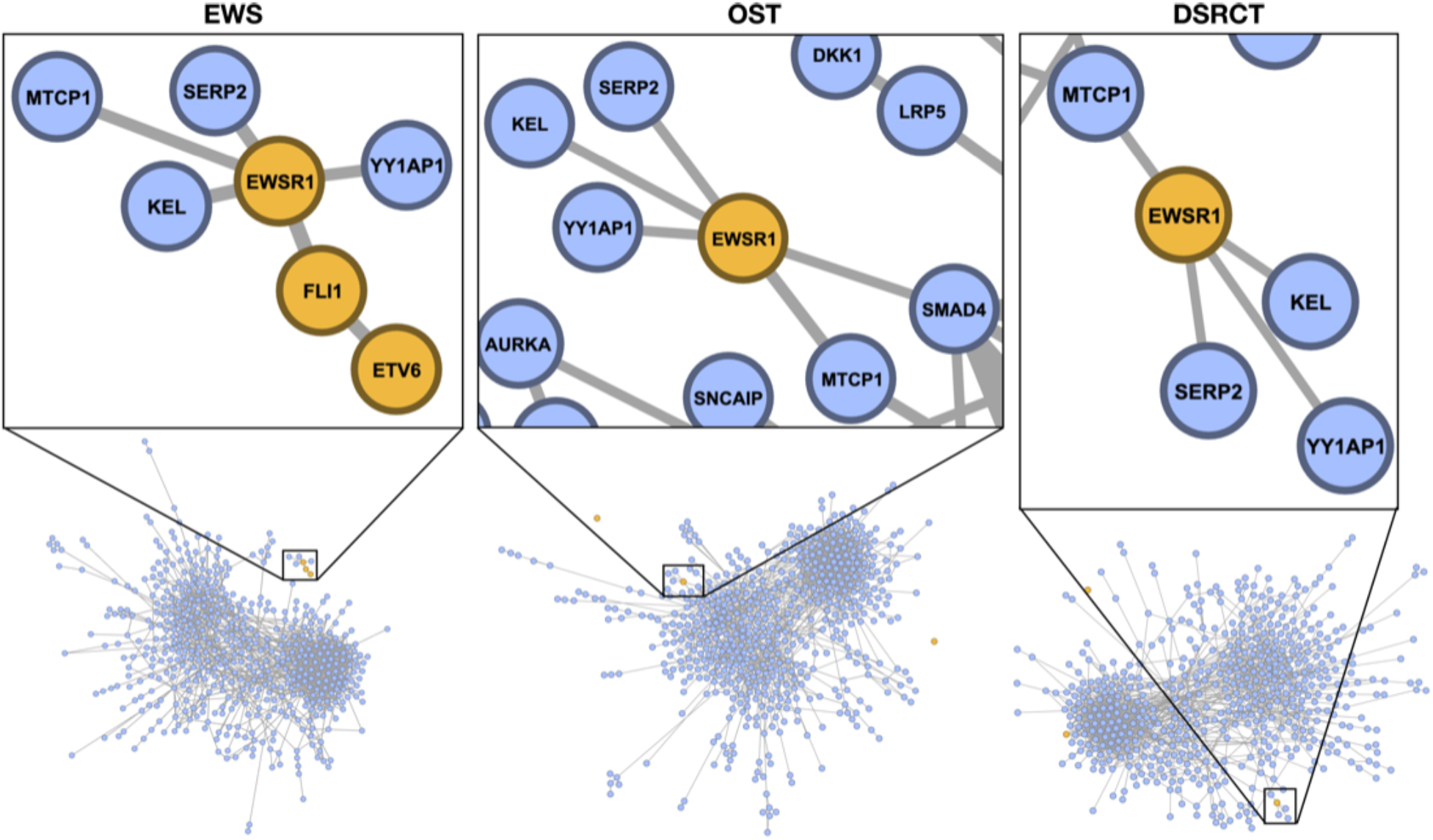
Critical curvature filtering of pediatric sarcoma networks. Functional community structures at the critical scale were realized by pruning bridges with negative critical curvature. The EWS network recovered the known functional *EWSR1-FLI1-ETV6* association.

#### 3.2.2 Multi-scale functional clustering: results

Hierarchical-acc clustering was performed on the EWS network 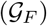. The resulting dendrogram encapsulated preferential gene clustering according to their *geometric cooperation.* As one would expect for EWS, *EWSR1, FLI1* and *ETV6* clustered together, highlighted in Figure 6. Importantly, this cluster was recovered in a purely agnostic fashion that is unique to the EWS network. As a side note, *KEL* and *SERP2* are leaf nodes attached to *EWSR1* in the original graph so it is not surprising that they are found in the same cluster. However, even this dependency is found to be less functionally relevant than the *EWSR1-FLI-ETV6* association, as demonstrated by the hierarchical ordering. Since the *EWSR1-FLI1* fusion has proven difficult to directly target [18], we investigated how they are affected by other interactions in the network, described in the next section.

**Figure 6:**
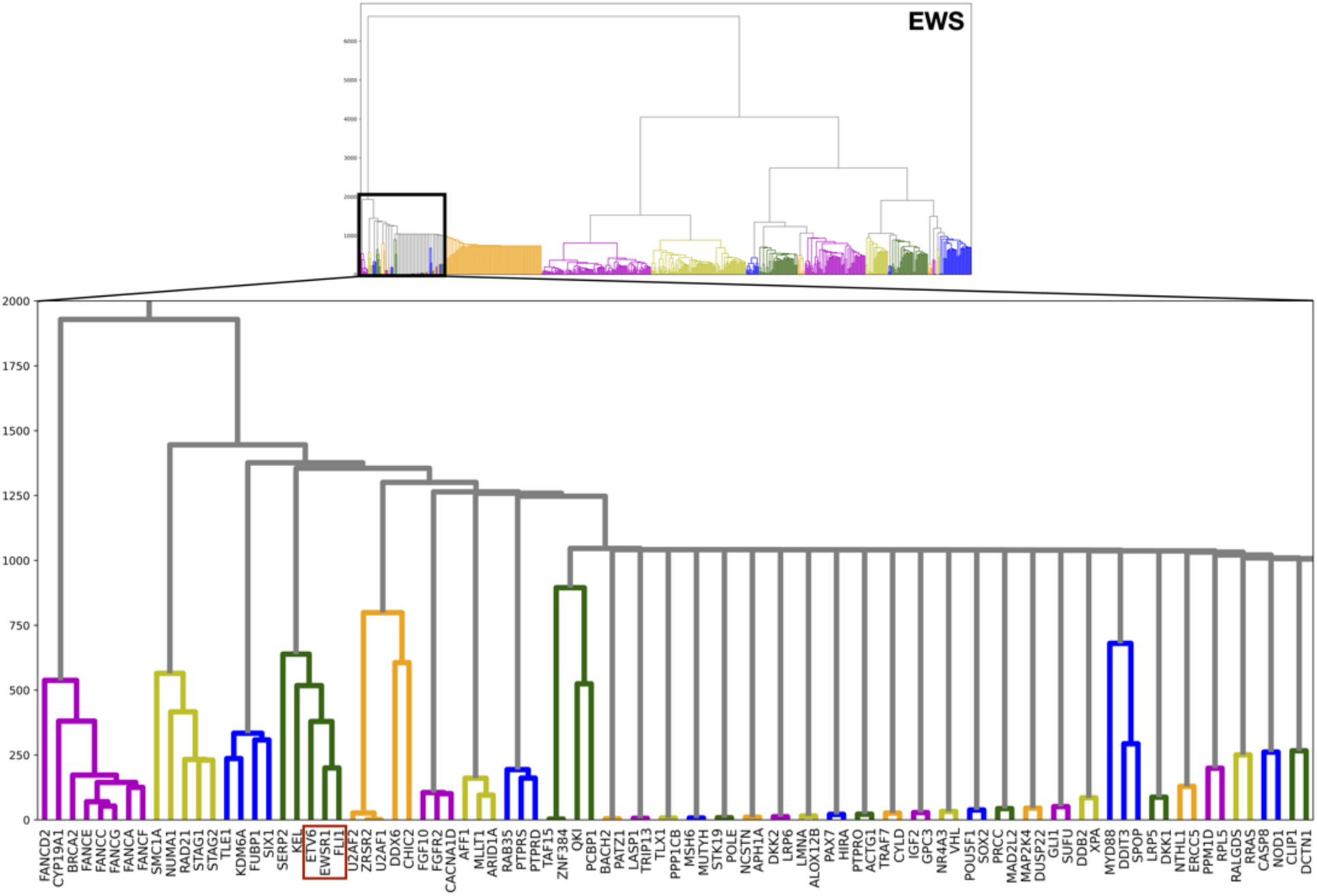
Hierarchical-acc clustering of the EWS network, highlighting the *EWSR1-FLI1-ETV6* association.

#### 3.2.3 Perturbation simulations: results

Perturbation simulations were performed on each edge in the network and curvature was computed for gene pairs *EWSR1-FLI1, FLI1-ETV6* and *EWSR1-ETV6* as described in Section 2.4.7 to assess the functional effect of targeted disruption in direct and indirect cooperation on the system.

The *net* change in curvature Δ between two nodes *r*, 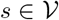 in response to perturbing edge 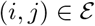 measures the net change in robustness, which is quantified as the difference in curvature after (i.e., with perturbed edge weight 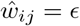) and before (i.e., baseline) effectively removing communication along the perturbed edge. The sign of Δ allows us to distinguish between *strengthening* and *weakening* effects, characterized respectively by an increase or decrease in curvature with respect to the baseline. Edges that disconnect the network when removed were eliminated from consideration because the distance between nodes linked by that edge approaches infinity as the edge is perturbed, essentially breaking the communication altogether. We then ranked the effects perturbing the remaining edges had on the *EWSR1-FLI1, FLI1-ETV6* and *EWSR1-ETV6* interactions. Perturbed edges with absolute value of effect greater than 1 x 10^-4^ (i.e., |Δ| > 1 x 10^-4^) on the *EWSR1-FLI1, FLI1-ETV6* and *EWSR1-ETV6* interactions are listed respectively in Tables 1, 2 and 3 along with the net effect Δ.

**Table 1:** Perturbed edges with the largest net effect (Δ) on *EWSR1-FLI1*

| Effect | Perturbed Edge | $\Delta$ | Figure |
| --- | --- | --- | --- |
| Positive | *ERG-FLI1* | 0.13028 | S1 |
|  | *EWSR1-ERCC5* | 0.02809 | S2 |
|  | *EWSR1-PCBP1* | 0.02442 | S3 |
| Negative | *JUN-ERG* | -0.09351 | S4 |
|  | *AR-POU5F1* | -0.04674 | S5 |
|  | *JUN-AR* | -0.02469 | S6 |
|  | *SMAD4-EWSR1* | -0.02456 | S7 |
|  | *EWSR1-TAF1* | -0.02209 | S8 |
|  | *BTK-EWSR1* | -0.02155 | S17 |
|  | *EWSR1-POU5F1* | -0.02109 | S16 |
|  | *EWSR1-FLI1* | -0.01803 |  |
|  | *EWSR1-HMGA1* | -0.01766 | S19 |
|  | *JUN-HMGA1* | -0.01726 | S20 |
|  | *ETV6-FLI1* | -0.01086 | S9 |
|  | *JUN-TAF1* | -0.01042 |  |
|  | *BARD1-EWSR1* | -0.00893 | S21 |
|  | *AKT1-MTCP1* | -0.00746 |  |
|  | *CRKL-ETV6* | -0.00574 | S10 |

Abbreviations: $\Delta$ : net effect or net difference in curvature (curvature with perturbed edge weight $\epsilon$ - baseline curvature).

**Table 2:** Perturbed edges with the largest net effect (Δ) on *FLI1-ETV6*

| Effect | Perturbed Edge | $\Delta$ | Figure |
| --- | --- | --- | --- |
| Positive | *ERG-FLI1* | 0.72147 | S1 |
|  | *ETV6-FLI1* | 0.70863 | S9 |
|  | *CRKL-ETV6* | 0.12042 | S10 |
|  | *ETV6-GAB2* | 0.08775 | S11 |
| Negative | *JUN-ERG* | -0.16018 | S4 |
|  | *STAT1-SYK* | -0.03720 | S12 |
|  | *JUN-STAT1* | -0.03720 | S13 |
|  | *SYK-GAB2* | -0.03720 |  |
|  | *SOS1-CRKL* | -0.02990 | S22 |
|  | *SOS1-ESR1* | -0.02990 | S23 |
|  | *JUN-SR1* | -0.02990 | S24 |
|  | *EWSR1-FLI1* | -0.00415 |  |
|  | *BTK-EWSR1* | -0.00164 | S17 |
|  | *BTK-WAS* | -0.00119 | S14 |
|  | *CRKL-WAS* | -0.00119 | S15 |

Abbreviations: $\Delta$ : net effect or net difference in curvature (curvature with perturbed edge weight $\epsilon$ - baseline curvature).

**Table 3:** Perturbed edges with the largest net effect (Δ) on *EWSR1-ETV6*

| Effect | Perturbed Edge | $\Delta$ | Figure |
| --- | --- | --- | --- |
| Positive | *EWSR1-ERCC5* | 0.02297 | S2 |
|  | *EWSR1-PCBP1* | 0.02144 | S3 |
|  | *BTK-WAS* | 0.00859 | S14 |
|  | *CRKL-WAS* | 0.00859 | S15 |
|  | *EWSR1-POU5F1* | 0.00383 | S16 |
| Negative | *ERG-FLI1* | -0.07069 | S1 |
|  | *ETV6-GAB2* | -0.05692 | S11 |
|  | *BTK-EWSR1* | -0.05339 | S17 |
|  | *SMAD4-EWSR1* | -0.03921 | S7 |
|  | *JUN-ERG* | -0.03168 | S4 |
|  | *AKT1-GAB2* | -0.02363 | S18 |
|  | *JUN-HMGA1* | -0.02218 | S20 |
|  | *AR-POU5F1* | -0.02191 |  |
|  | *MAPK1-GAB2* | -0.02154 | S25 |
|  | *RB1-TAF1* | -0.01824 | S26 |
|  | *EWSR1-TAF1* | -0.01785 | S8 |
|  | *EWSR1-HMGA1* | -0.01492 | S19 |
|  | *AKT1-AR* | -0.01082 |  |
|  | *RB1-MAPK1* | -0.01038 | S27 |
|  | *AKT1-MTCP1* | -0.01030 |  |
|  | *CRKL-ETV6* | -0.01001 | S10 |
|  | *EWSR1-MTCP1* | -0.00938 | S28 |
|  | *MAPK1-SMAD4* | -0.00874 | S29 |
|  | *BARD1-EWSR1* | -0.00448 | S21 |
|  | *ETV6-FLI1* | -0.00414 | S9 |
|  | *EWSR1-FLI1* | -0.00394 |  |
|  | *BRCA1-BARD1* | -0.00275 | S31 |
|  | *HSP90AA1-NDRG1* | -0.00230 |  |
|  | *EWSR1-NDRG1* | -0.00223 | S30 |
|  | *MAPK1-HSP90AA1* | -0.00139 |  |
|  | *CREBBP-EWSR1* | -0.00125 |  |
|  | *BRCA1-AKT1* | -0.00121 |  |
|  | *ERG-SETDB1* | -0.00021 | S32 |
|  | *BARD1-SETDB1* | -0.00021 | S33 |
|  | *MAPK1-NCOA1* | -0.00010 | S34 |
|  | *CREBBP-NCOA1* | -0.00010 |  |

Abbreviations: $\Delta$ : net effect or net difference in curvature (curvature with perturbed edge weight $\epsilon$ - baseline curvature).

To complement these tables and provide additional information on the network response to the simulated disruption, we looked at other *EWSR1/FLI1/ETV6-interactions* affected by the perturbed edges listed in Tables 1, 2 and 3, provided in the Supplemental Material. Barplots of the net change in curvature (Δ), where red represents a decrease in robustness and blue represents an increase, and line-plots of the curvature as a function of the decreasing perturbed edge weight are plotted for affected interactions with absolute value of effect greater than 0.2 (|Δ| > 0.2). The curvature for three additional perturbed edge-weights near 0 are shown to highlight trends as the edge is virtually cut. The figure reference corresponding to each perturbed edge with at least one absolute effect greater than 0.2 are provided in Tables 1, 2 and 3.

## 4 Discussion

In this work, we utilized a network version of the geometric concept of curvature to model information variability, robustness, and dysregulation of cancer gene networks. Pediatric sarcomas (PS) represent a phenotypically diverse group of malignant solid tumors [2]. A subset of PS is characterized by oncogenic driver fusion genes such as EWS-FLI1 in Ewing sarcoma, EWS-WT1 in desmoplastic small round cell tumor (DSRCT), and PAX3/7-FOXO1 in fusion-positive rhabdomyosarcoma [20]. Given the heterogeneous nature of PS and often overlapping microscopic structural features (histology) across different PS subtypes, the presence and detection of driver fusion genes in PS has aided in the diagnostic classification of these tumors. Here, we demonstrate that analysis of the curvature within GEPs as a function of scale is able to define robust networks that distinguish subtypes of PS. These approaches may therefore have utility as a RNA-based classifier aiding the diagnosis of PS subtypes.

Given the lack of other driver mutations that typify the mutational landscape of PS, or pediatric tumors in general, direct targeting of fusion oncogenes has seemed a logical strategy for treating fusion-positive PS [18]. However, development of drugs that can selectively target and inhibit the activity of fusion oncogenes has remained elusive [18]. Therefore, development of strategies that identify targets that indirectly disrupt the key functional interactions nucleated by “undruggable” fusion oncoproteins, or enable the identification of driver mutations amidst a low tumor mutational landscape characteristic of pediatric cancers [12], addresses a critical unmet need in pediatric oncology.

The work presented provides a novel approach for mining genomic sequencing data to aid diagnostic classification of PS and identify potential therapeutic targets not readily assessable by merely cataloguing a tumor’s set of mutations. Systematic selective targeting of genes involved in critical interactions (e.g., *EWSR1-FLI1-ETV6* interaction) and the functional consequences of inhibiting critical interactions in *in vivo* tumor models of PS will provide future validation of this approach and inform future applications of curvature analysis in pediatric oncology.

## Code availability

The code written in Python is available upon request.

## Acknowledgements

This research was funded in part by grants from the Air Force Office of Scientific Research (FA9550-17-1-0435, FA9550-20-1-0029), NIH grants (R01-AG048769, R21-CA234752), MSK Cancer Center Support Grant/Core Grant (P30 CA008748), and a grant from Breast Cancer Research Foundation (BCRF-17-193).

## Author Contributions

R.E. and A.T. developed the mathematical methods, and J.O. developed the bioinformatic analysis. A.L.K. and F.D.C. conceived the project and provided key biological and clinical analysis and interpretation. J.D. provided insights into interpreting the results and clarifying the technical methods. R.E. wrote the paper, and all authors edited the paper.

## References

[1] Luigi Ambrosio. Lecture notes on optimal transport problems. In Mathematical Aspects of Evolving Interfaces, pages 1–52. Springer, 2003.

[2] Jennifer L Anderson, Christopher T Denny, William D Tap, and Noah Federman. Pediatric sarcomas: translating molecular pathogenesis of disease to novel therapeutic possibilities. Pediatric research, 72(2):112–121, 2012.

[3] Matthew R Avenarius, Cecelia R Miller, Michael A Arnold, Selene Koo, Ryan Roberts, Martin Hobby, Thomas Grossman, Yvonne Moyer, Richard K Wilson, Elaine R Mardis, et al. Genetic characterization of pediatric sarcomas by targeted rna sequencing. The Journal of Molecular Diagnostics, 22(10):1238–1245, 2020.

[4] D Bakry and M Émery. Diffusions hypercontractives séminaire de probabilités, xix. Lecture Notes in Mathematics, 1123:177–206, 1985.

[5] Frank Bauer, Jürgen Jost, and Shiping Liu. Ollivier-Ricci curvature and the spectrum of the normalized graph laplace operator. Math. Res. Lett., 19:1185–1205, 2012.

[6] Kévin Bourcier, Axel Le Cesne, Lambros Tselikas, Julien Adam, Olivier Mir, Charles Honore, and Thierry de Baere. Basic knowledge in soft tissue sarcoma. Cardiovascular and interventional radiology, 42(9):1255–1261, 2019.

[7] Manfredo Perdigao do Carmo. Riemannian Geometry. Birkhäuser, 1992.

[8] Debyani Chakravarty, Jianjiong Gao, Sarah Phillips, Ritika Kundra, Hongxin Zhang, Jiaojiao Wang, Julia E Rudolph, Rona Yaeger, Tara Soumerai, Moriah H Nissan, et al. Oncokb: a precision oncology knowledge base. JCO precision oncology, (1):1–16, 2017.

[9] Hamza Farooq, Yongxin Chen, Tryphon T Georgiou, Allen Tannenbaum, and Christophe Lenglet. Network curvature as a hallmark of brain structural connectivity. Nature Communications, 10(1):1–11, 2019.

[10] Robin Forman. Bochner’s method for cell complexes and combinatorial ricci curvature. Discrete Comput Geometry, 29:323–374, 2003.

[11] Adam Gosztolai and Alexis Arnaudon. Unfolding the multiscale structure of networks with dynamical ollivier-ricci curvature. https://arxiv.org/abs/2106.05847, 2021.

[12] Susanne N Gröbner, Barbara C Worst, Joachim Weischenfeldt, Ivo Buchhalter, Kortine Kleinheinz, Vasilisa A Rudneva, Pascal D Johann, Gnana Prakash Balasubramanian, Maia Segura-Wang, Sebastian Brabetz, et al. The landscape of genomic alterations across childhood cancers. Nature, 555(7696):321–327, 2018.

[13] Chiang-Ching Huang, Colleen Cutcliffe, Cheryl Coffin, Poul HB Sorensen, J Bruce Beckwith, and Elizabeth J Perlman. Classification of malignant pediatric renal tumors by gene expression. Pediatric blood & cancer, 46(7):728–738, 2006.

[14] Trey Ideker, Owen Ozier, Benno Schwikowski, and Andrew F Siegel. Discovering regulatory and signalling circuits in molecular interaction networks. Bioinformatics, 18(suppl_1):S233–S240, 2002.

[15] Michael S Isakoff, Robert Goldsby, Doojduen Villaluna, Mark D Krailo, Pooja Hingorani, Anderson Collier, Carol D Morris, E Anders Kolb, John J Doski, Richard B Womer, et al. A phase ii study of eribulin in recurrent or refractory osteosarcoma: a report from the children’s oncology group. Pediatric blood & cancer, 66(2):e27524, 2019.

[16] TS Keshava Prasad, Renu Goel, Kumaran Kandasamy, Shivakumar Keerthikumar, Sameer Kumar, Suresh Mathivanan, Deepthi Telikicherla, Rajesh Raju, Beema Shafreen, Abhilash Venugopal, et al. Human protein reference database?2009 update. Nucleic acids research, 37(suppl_1):D767–D772, 2009.

[17] Christian Koelsche, Daniel Schrimpf, Damian Stichel, Martin Sill, Felix Sahm, David E Reuss, Mirjam Blattner, Barbara Worst, Christoph E Heilig, Katja Beck, et al. Sarcoma classification by dna methylation profiling. Nature communications, 12(1):1–10, 2021.

[18] Heinrich Kovar. Blocking the road, stopping the engine or killing the driver? advances in targeting ews/fli-1 fusion in ewing sarcoma as novel therapy. Expert opinion on therapeutic targets, 18(11):1315–1328, 2014.

[19] John Lott and Cédric Villani. Ricci curvature for metric-measure spaces via optimal transport. Annals of Mathematics, pages 903–991, 2009.

[20] Crystal L Mackall, Paul S Meltzer, and Lee J Helman. Focus on sarcomas. Cancer cell, 2(3):175–178, 2002.

[21] Isabella WY Mak, Nathan Evaniew, and Michelle Ghert. Lost in translation: animal models and clinical trials in cancer treatment. American journal of translational research, 6(2):114, 2014.

[22] Suman Malempati, Brenda Weigel, Ashish M Ingle, Charlotte H Ahern, Julie M Carroll, Charles T Roberts, Joel M Reid, Stephen Schmechel, Stephan D Voss, Steven Y Cho, et al. Phase i/ii trial and pharmacokinetic study of cixutumumab in pediatric patients with refractory solid tumors and ewing sarcoma: a report from the children’s oncology group. Journal of Clinical Oncology, 30(3):256, 2012.

[23] Yann Ollivier. Ricci curvature of markov chains on metric spaces. J. Functional Analysis, 256:810–864, 2009.

[24] Alberto S Pappo, Gilles Vassal, John J Crowley, Vanessa Bolejack, Pancras CW Hogendoorn, Rashmi Chugh, Marc Ladanyi, Joseph F Grippo, Georgina Dall, Arthur P Staddon, et al. A phase 2 trial of r1507, a monoclonal antibody to the insulin-like growth factor-1 receptor (igf-1r), in patients with recurrent or refractory rhabdomyosarcoma, osteosarcoma, synovial sarcoma, and other soft tissue sarcomas: Results of a sarcoma alliance for research through collaboration study. Cancer, 120(16):2448–2456, 2014.

[25] Suraj Peri, J Daniel Navarro, Ramars Amanchy, Troels Z Kristiansen, Chandra Kiran Jonnalagadda, Vineeth Surendranath, Vidya Niranjan, Babylakshmi Muthusamy, TKB Gandhi, Mads Gronborg, et al. Development of human protein reference database as an initial platform for approaching systems biology in humans. Genome research, 13(10):2363–2371, 2003.

[26] Maryam Pouryahya, Jung Hun Oh, Pedram Javanmard, James C Mathews, Zehor Belkhatir, Joseph O Deasy, and Allen Tannenbaum. awcluster: A novel integrative network-based clustering of multiomics for subtype analysis of cancer data. IEEE/ACM Transactions on Computational Biology and Bioinformatics, 2020.

[27] Romeil Sandhu et al. Graph curvature for differentiating cancer networks. Scientific Reports, 5(1):1–13, 2015.

[28] Inga-Marie Schaefer, Gregory M Cote, and Jason L Hornick. Contemporary sarcoma diagnosis, genetics, and genomics. Journal of Clinical Oncology, 36(2):101–110, 2018.

[29] Eric S Schafer, Rachel E Rau, Stacey L Berg, Xiaowei Liu, Charles G Minard, Alexander JR Bishop, J Carolina Romero, M John Hicks, Marvin D Nelson Jr, Stephan Voss, et al. Phase 1/2 trial of talazoparib in combination with temozolomide in children and adolescents with refractory/recurrent solid tumors including ewing sarcoma: A children’s oncology group phase 1 consortium study (advl1411). Pediatric blood & cancer, 67(2):e28073, 2020.

[30] Cédric Villani. Topics in Optimal Transportation. Number 58. American Mathematical Soc., 2003.

[31] Cédric Villani. Optimal Transport: Old and New, volume 338. Springer Science & Business Media, 2008.

[32] Max-K von Renesse and Karl-Theodor Sturm. Transport inequalities, gradient estimates, entropy and ricci curvature. Communications on Pure and Applied Mathematics, 58(7):923–940, 2005.

[33] Anne B Warwick, Suman Malempati, Mark Krailo, Allen Melemed, Richard Gorlick, Matthew M Ames, Stephanie L Safgren, Peter C Adamson, and Susan M Blaney. Phase 2 trial of pemetrexed in children and adolescents with refractory solid tumors: a children’s oncology group study. Pediatric blood & cancer, 60(2):237–241, 2013.

[34] Brenda Weigel, Suman Malempati, Joel M Reid, Stephan D Voss, Steven Y Cho, Helen X Chen, Mark Krailo, Doojduen Villaluna, Peter C Adamson, and Susan M Blaney. Phase 2 trial of cixutumumab in children, adolescents, and young adults with refractory solid tumors: a report from the children’s oncology group. Pediatric blood & cancer, 61(3):452–456, 2014.

[35] S Peter Wu, Benjamin T Cooper, Fang Bu, Christopher J Bowman, J Keith Killian, Jonathan Serrano, Shiyang Wang, Twana M Jackson, Daniel Gorovets, Neerav Shukla, et al. Dna methylation–based classifier for accurate molecular diagnosis of bone sarcomas. JCO precision oncology, 1:1–11, 2017.

