## Supplementary Material for "Dynamic Network Curvature Analysis of RNA-Seq Data in Sarcoma"

### Supplemental Material

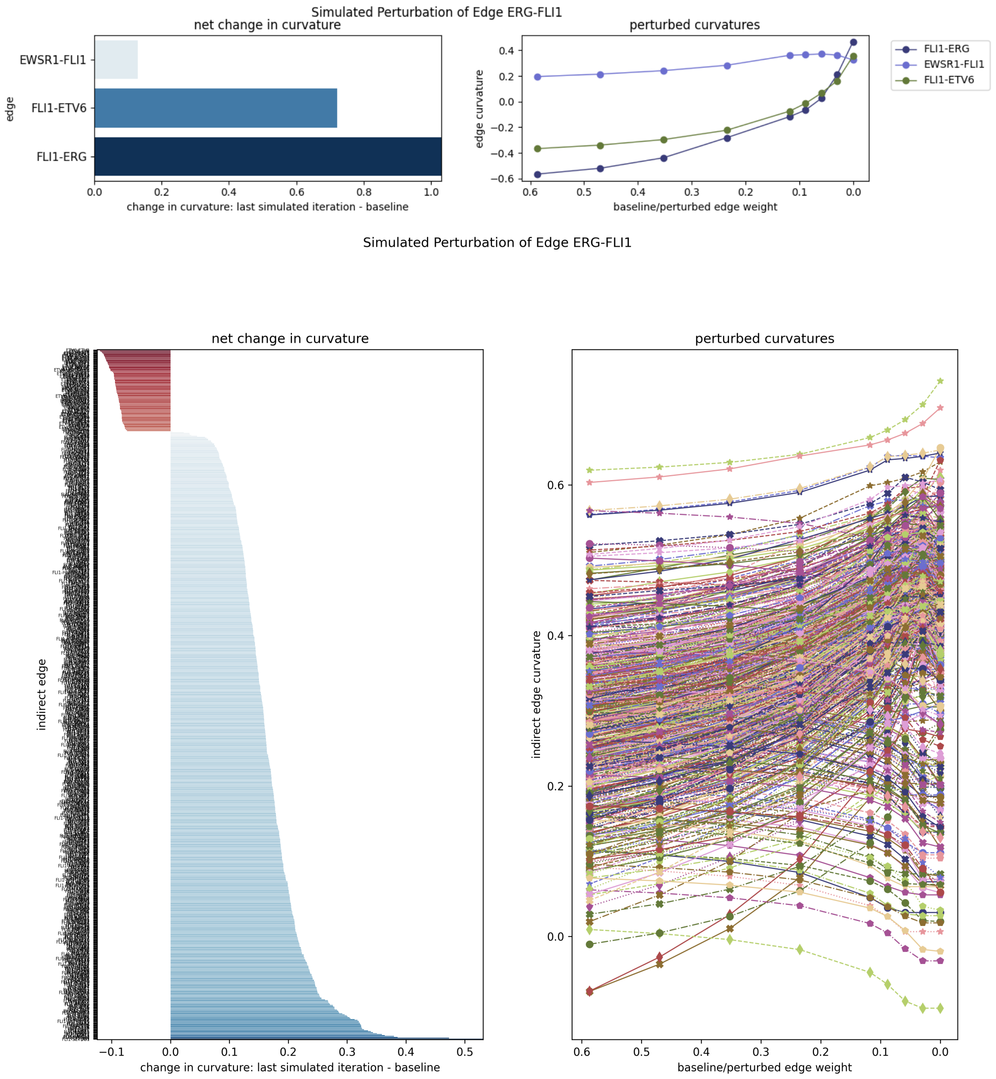


Figure S1: Perturbing edge *ERG-FLI1*. (Top) Direct affected interactions, i.e., edges. (Bottom) Indirect affected interactions.

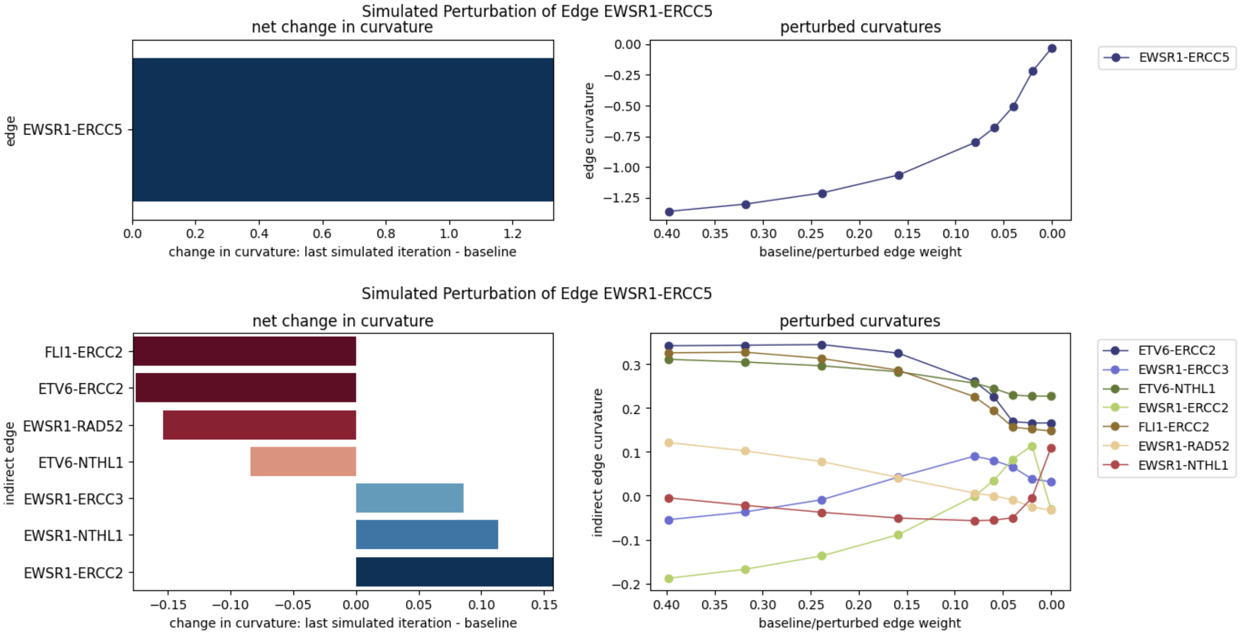


Figure S2: Perturbing edge *EWSR1-ERCC5*. (Top) Direct affected interactions, i.e., edges. (Bottom) Indirect affected interactions.

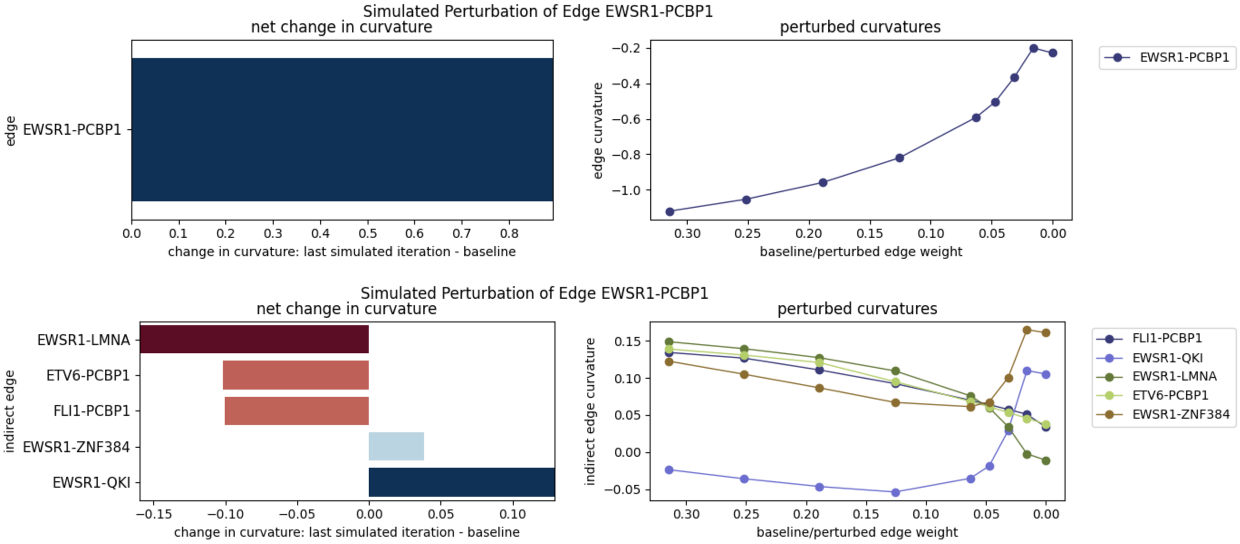


Figure S3: Perturbing edge *EWSR1-PCBP1*. (Top) Direct affected interactions, i.e., edges. (Bottom) Indirect affected interactions.

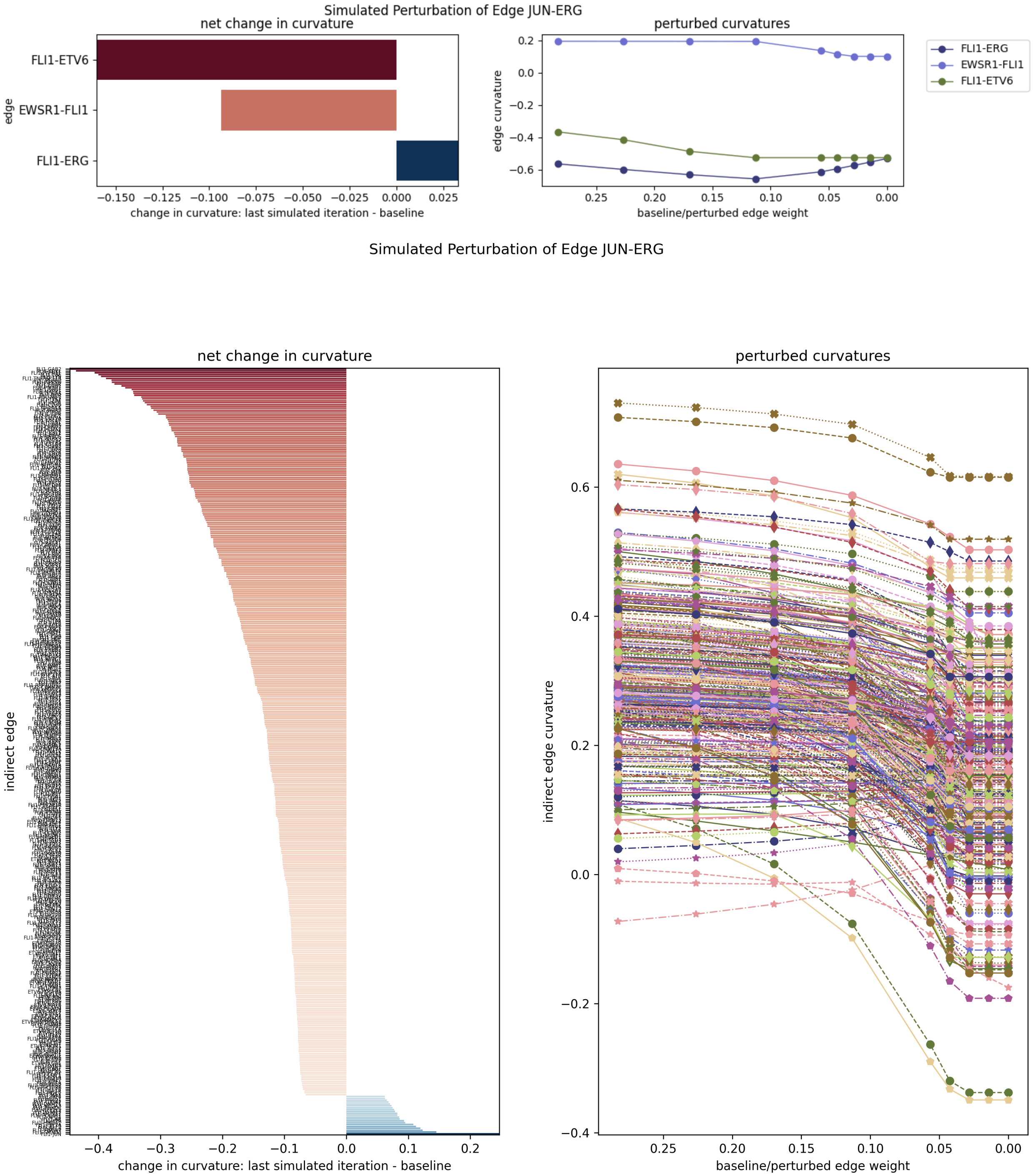


Figure S4: Perturbing edge *JUN-ERG*. (Top) Direct affected interactions, i.e., edges. (Bottom) Indirect affected interactions.

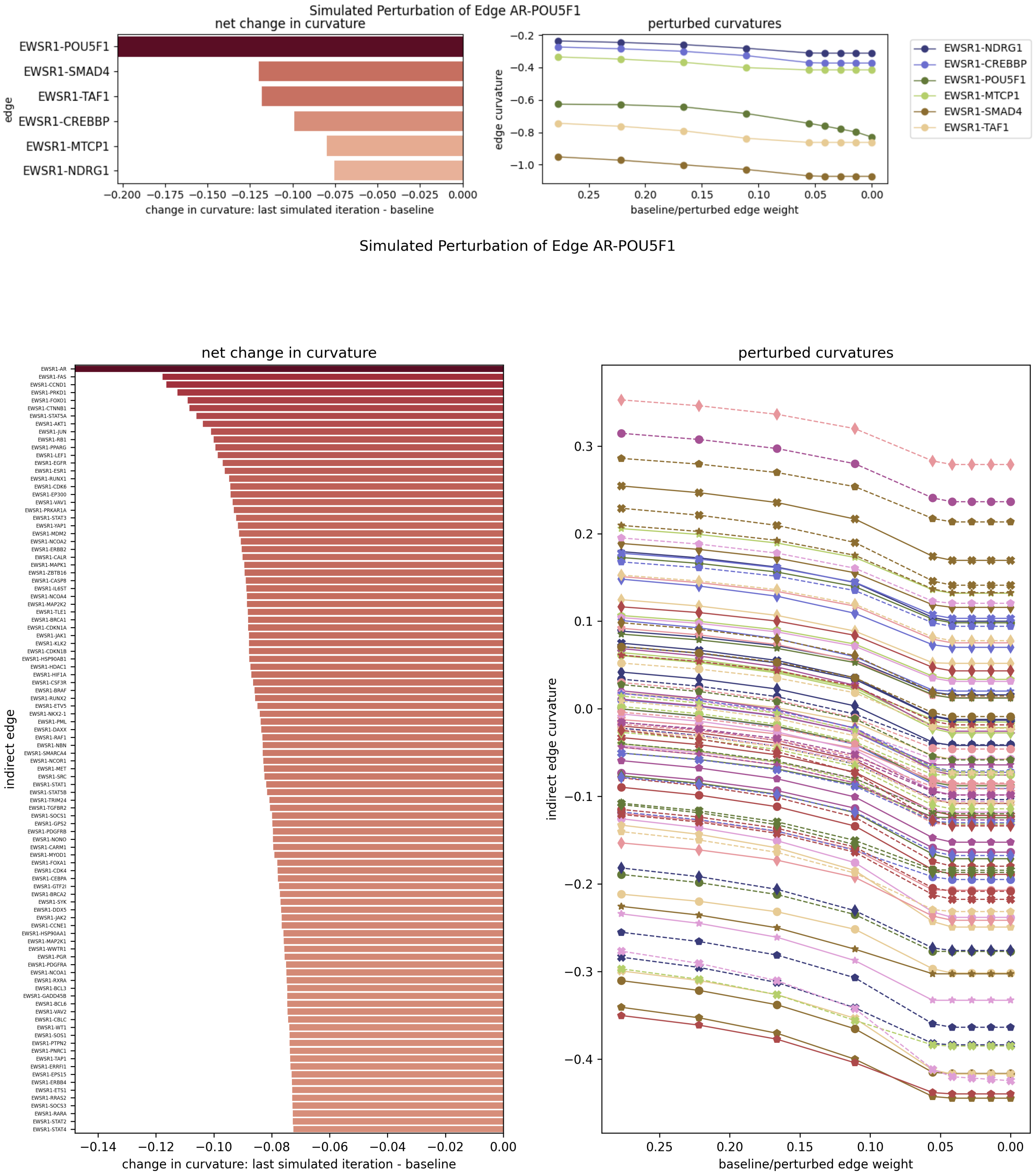


Figure S5: Perturbing edge *AR-POU5F1*. (Top) Direct affected interactions, i.e., edges. (Bottom) Indirect affected interactions.

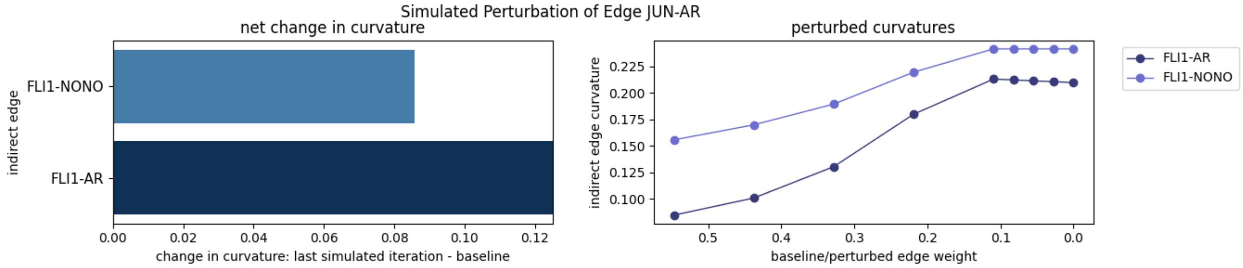


Figure S6: Perturbing edge $JUN$-$AR$.

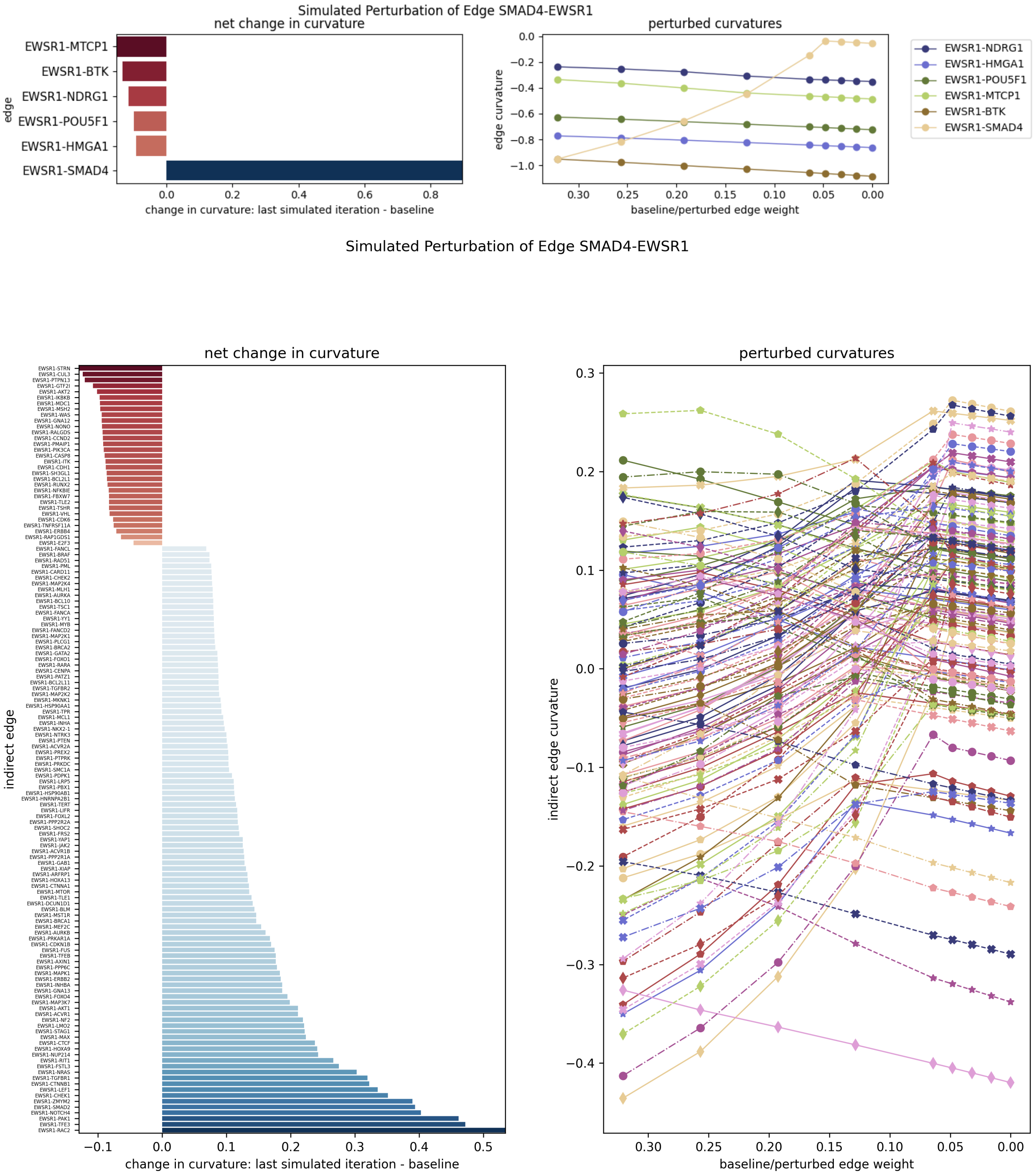


Figure S7: Perturbing edge *SMAD4-EWSR1*. (Top) Direct affected interactions, i.e., edges. (Bottom) Indirect affected interactions.

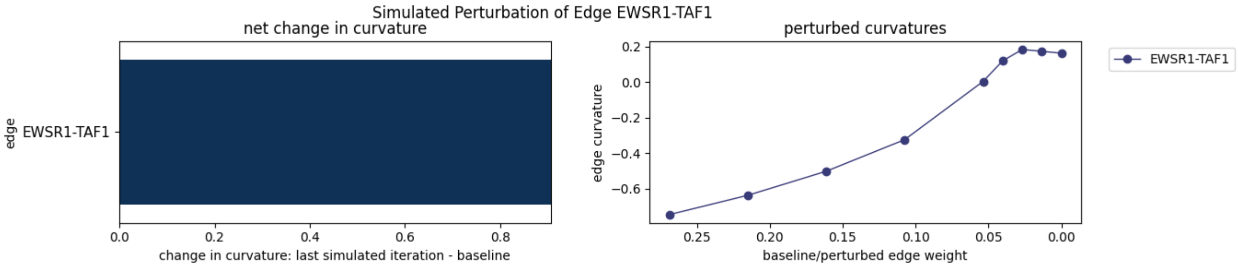


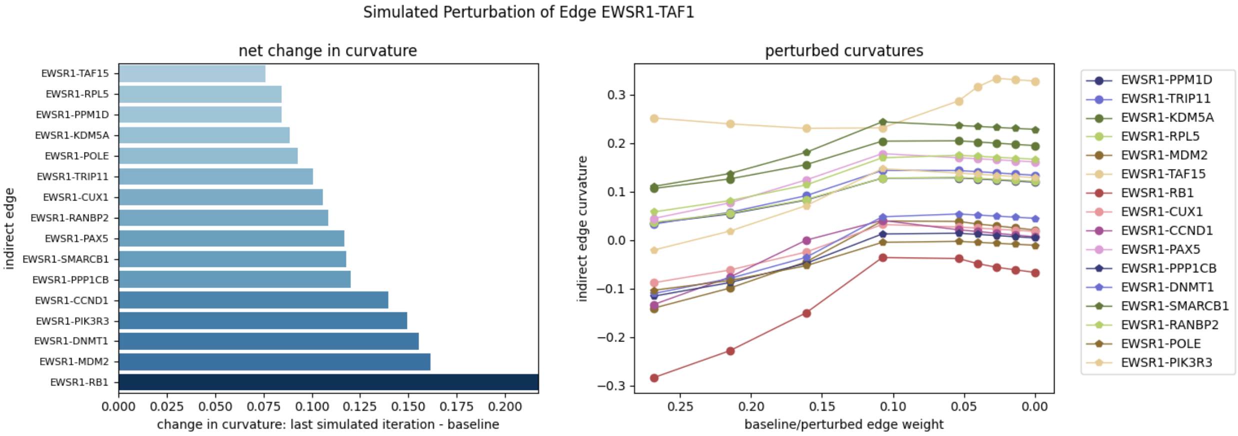


Figure S8: Perturbing edge *EWSR1-TAF1*. (Top) Direct affected interactions, i.e., edges. (Bottom) Indirect affected interactions.

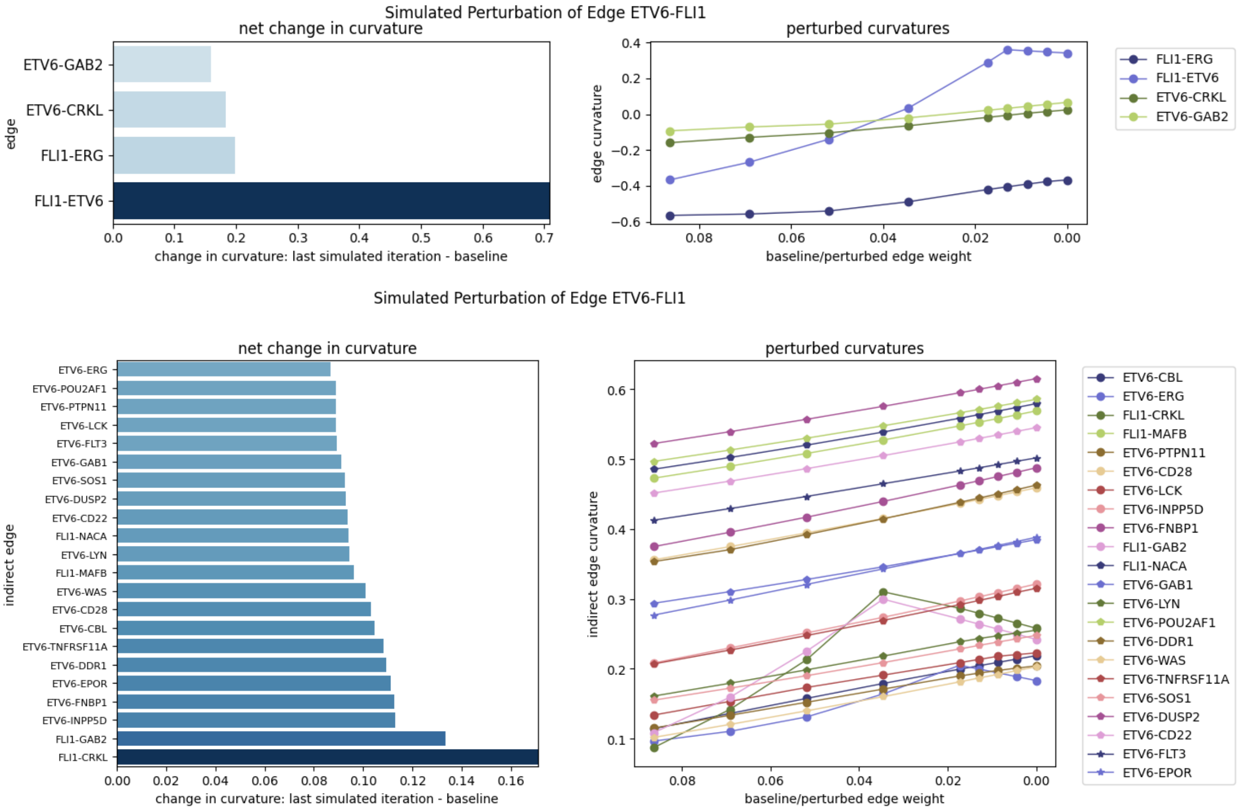


Figure S9: Perturbing edge *ETV6-FLI1*. (Top) Direct affected interactions, i.e., edges. (Bottom) Indirect affected interactions.

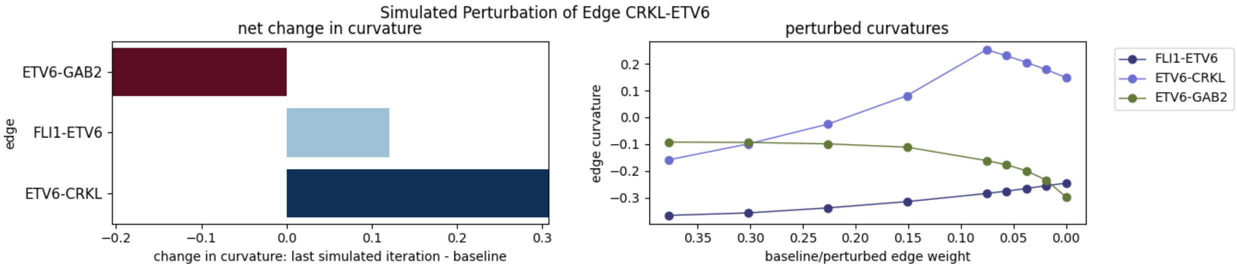


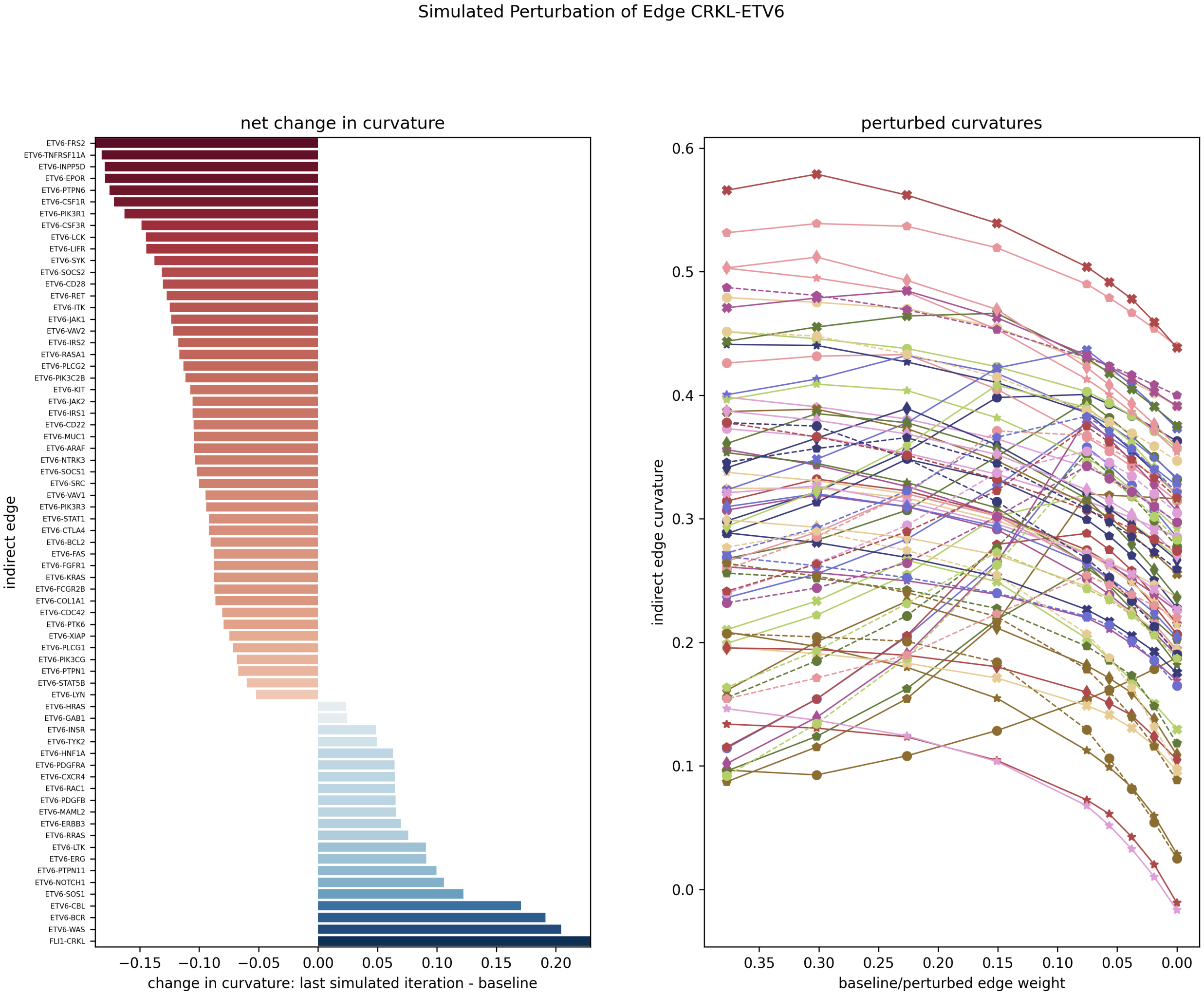


Figure S10: Perturbing edge *CRKL-ETV6*. (Top) Direct affected interactions, i.e., edges. (Bottom) Indirect affected interactions.

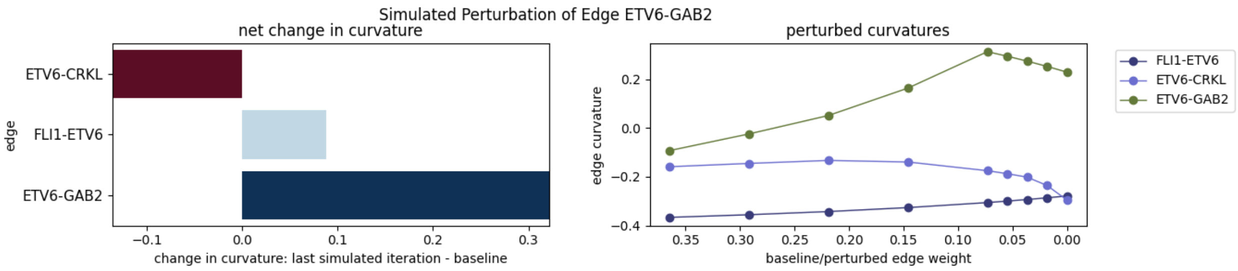


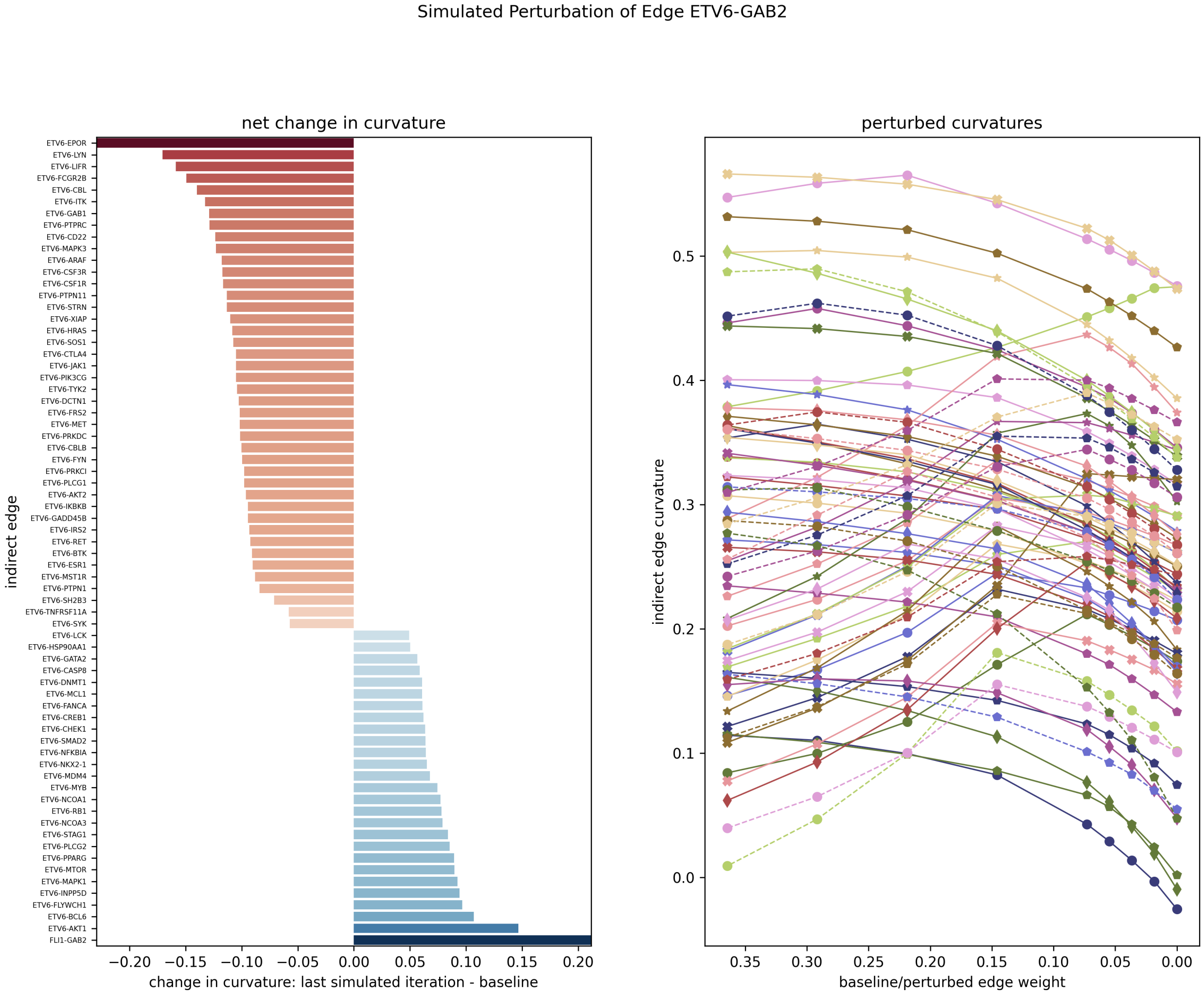


Figure S11: Perturbing edge *ETV6-GAB2*. (Top) Direct affected interactions, i.e., edges. (Bottom) Indirect affected interactions.

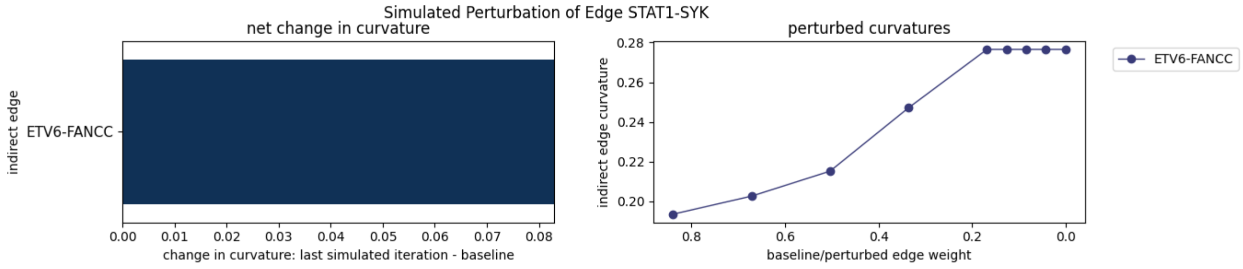


Figure S12: Perturbing edge *STAT1-SYK*.

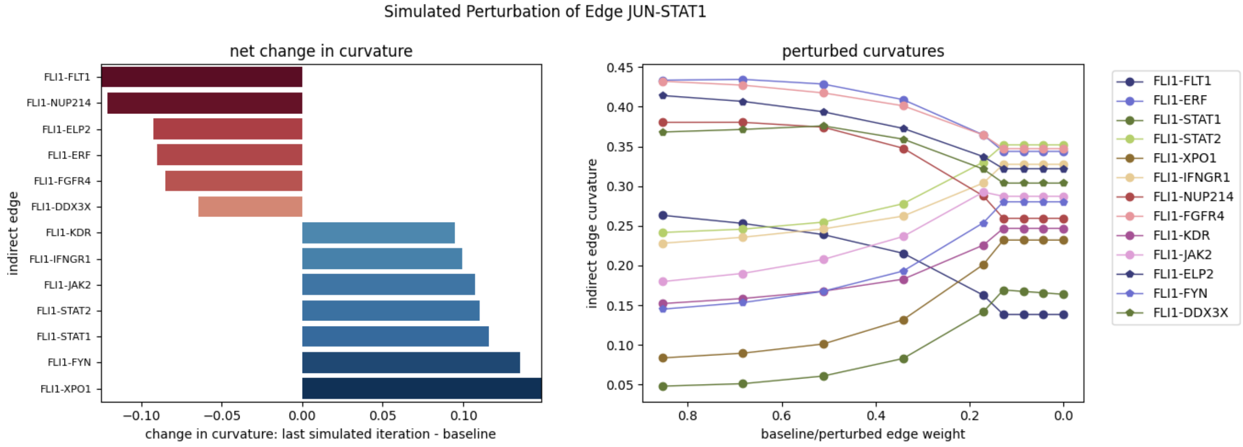


Figure S13: Perturbing edge *JUN-STAT1*.

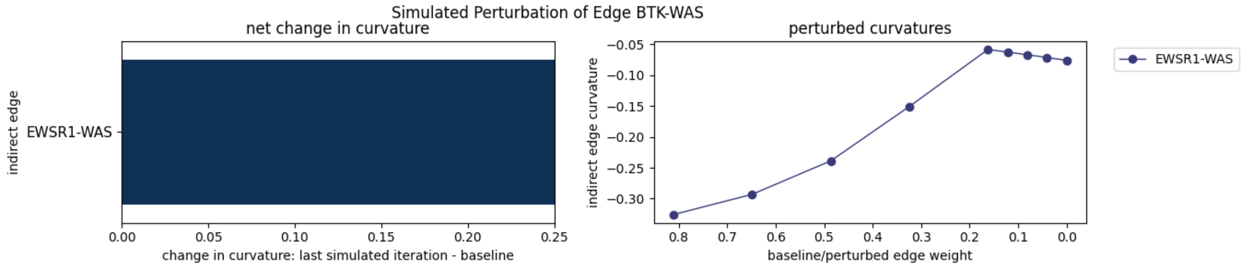


Figure S14: Perturbing edge *BTK-WAS*.

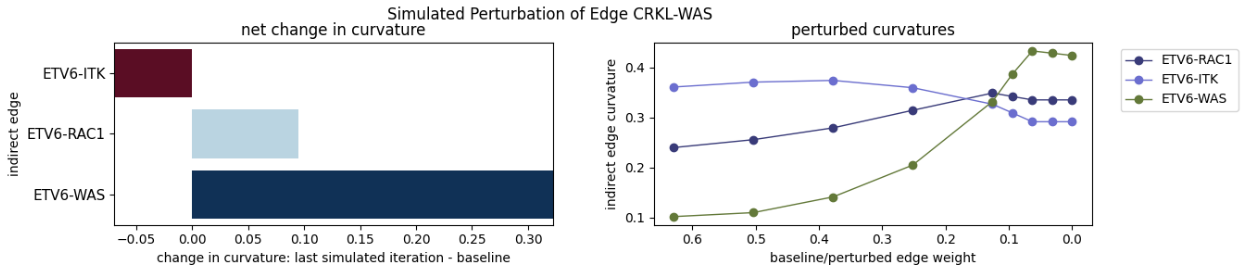


Figure S15: Perturbing edge *CRKL-WAS*.

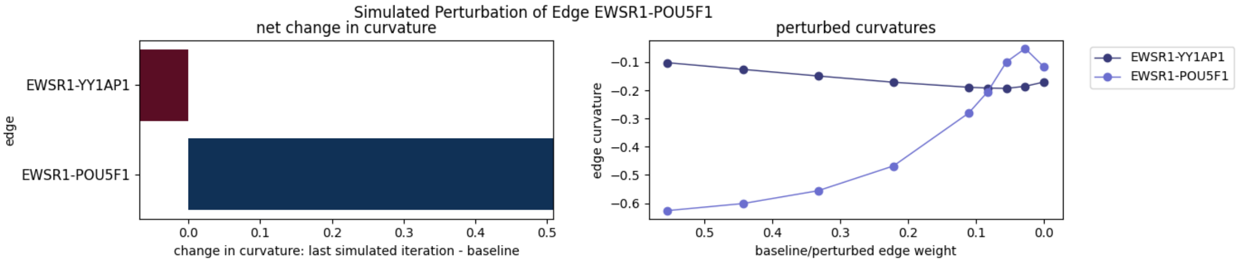


Figure S16: Directly affected interactions by perturbing edge *EWSR1-POU5F1*.

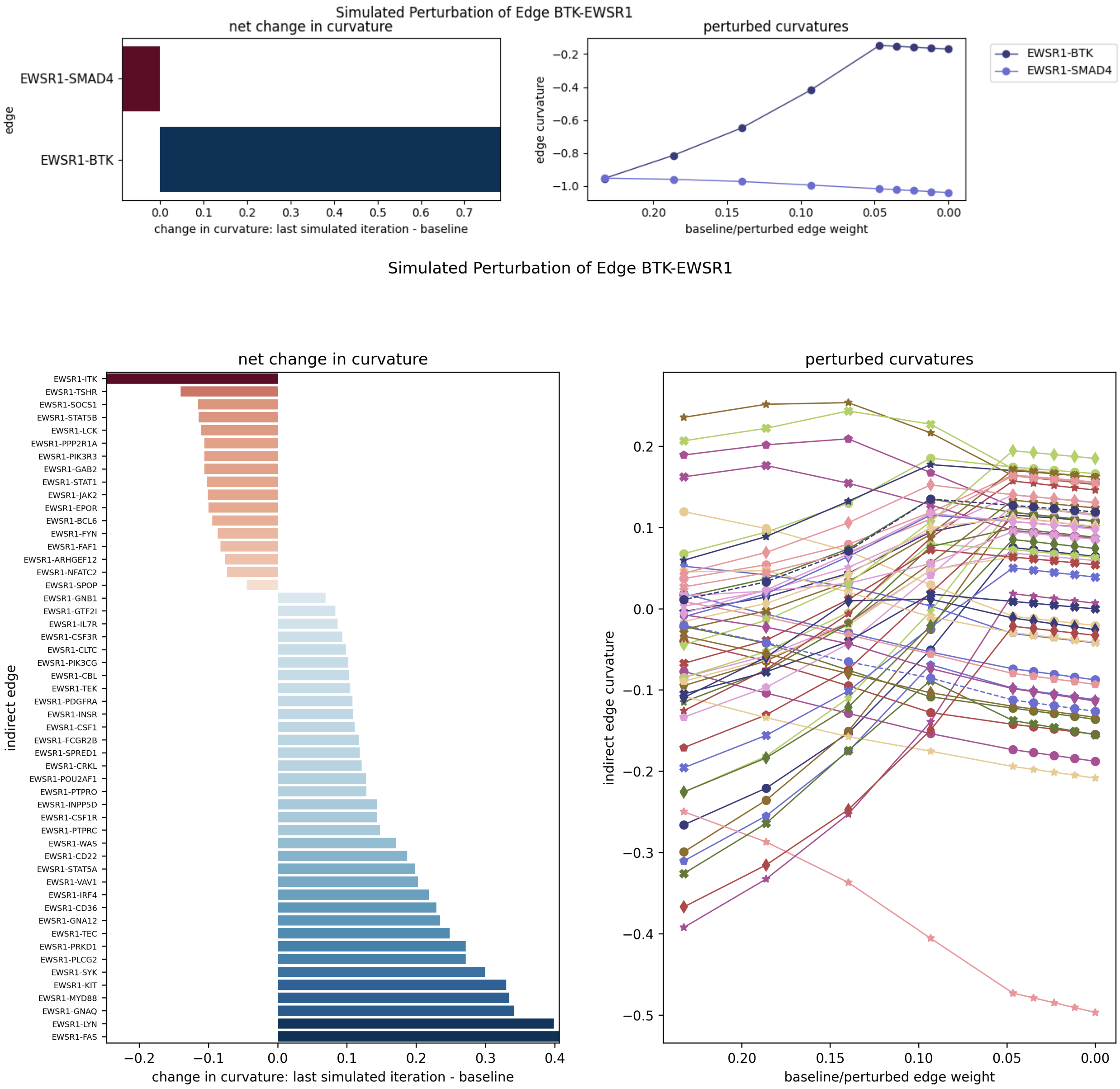


Figure S17: Perturbing edge *BTK-EWRS1*. (Top) Direct affected interactions, i.e., edges. (Bottom) Indirect affected interactions.

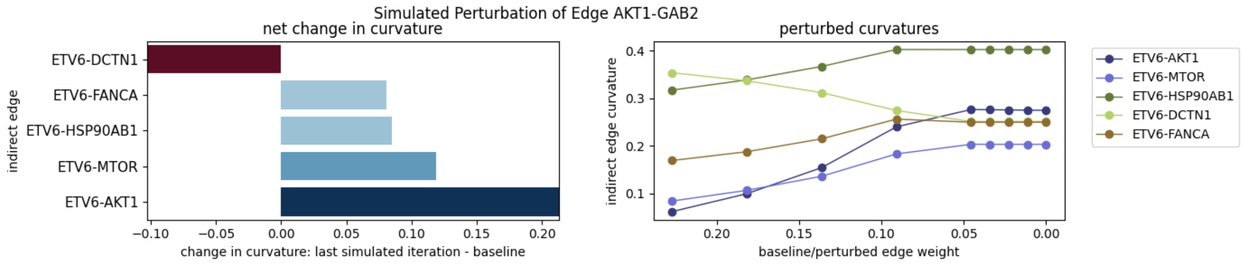


Figure S18: Perturbing edge *AKT1-GAB2*.

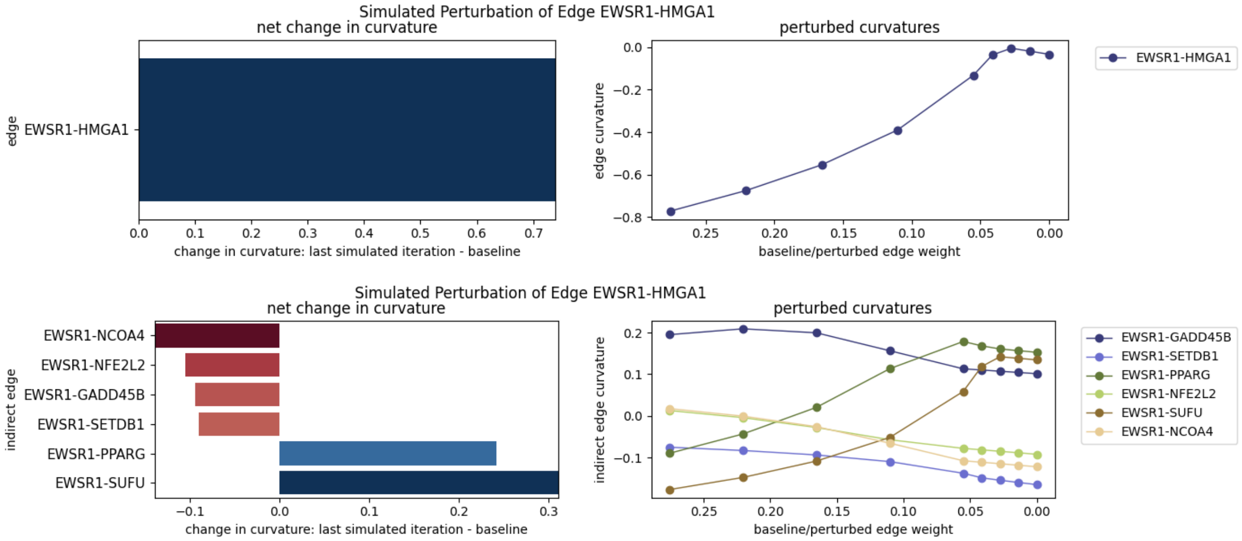


Figure S19: Perturbing edge *EWSR1-HMGA1*. (Top) Direct affected interactions, i.e., edges. (Bottom) Indirect affected interactions.

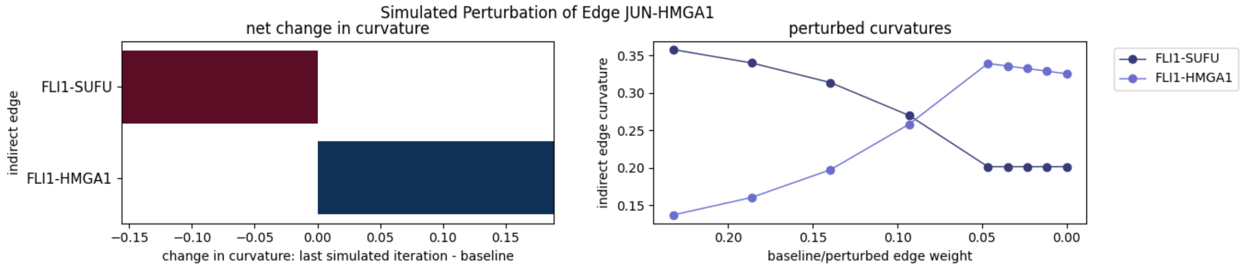


Figure S20: Perturbing edge *JUN-HMGA1*.

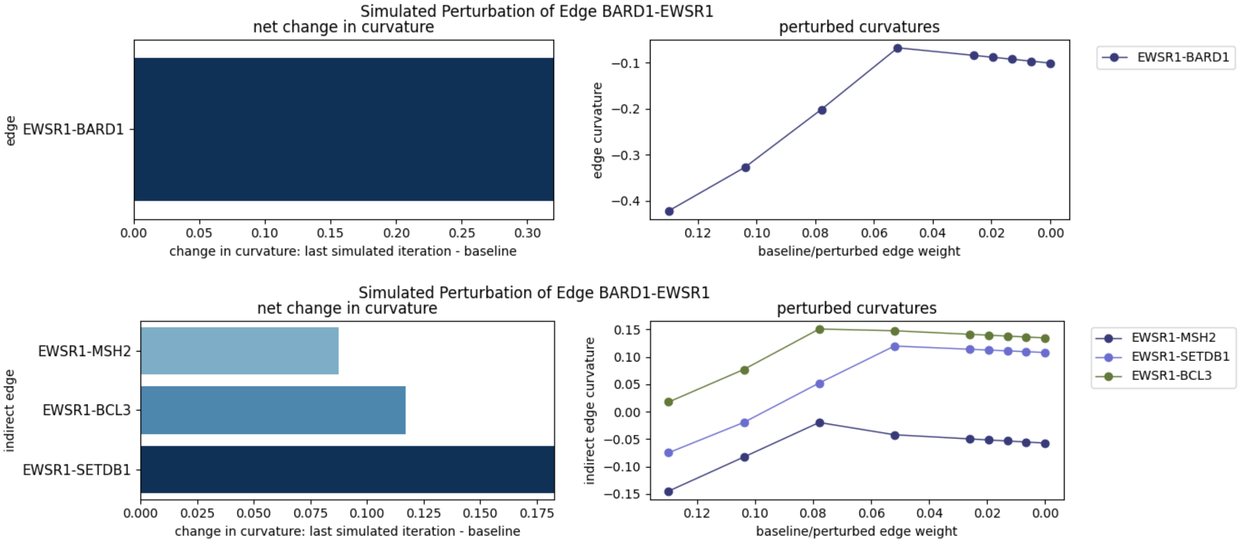


Figure S21: Perturbing edge *BARD1-EWSR1*. (Top) Direct affected interactions, i.e., edges. (Bottom) Indirect affected interactions.

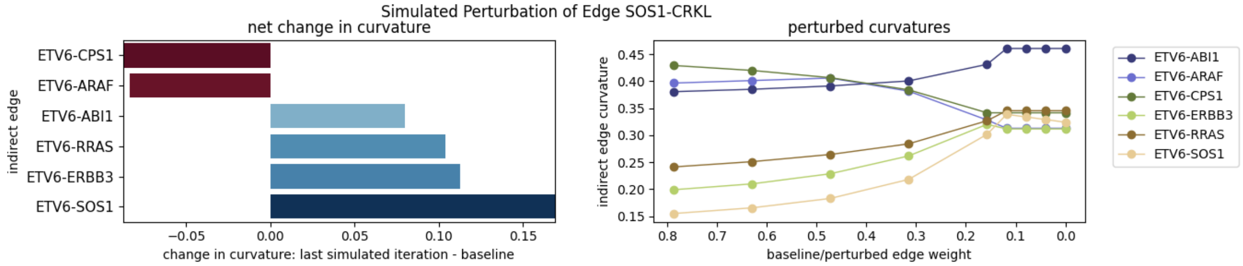


Figure S22: Perturbing edge *SOS1-CRKL*.

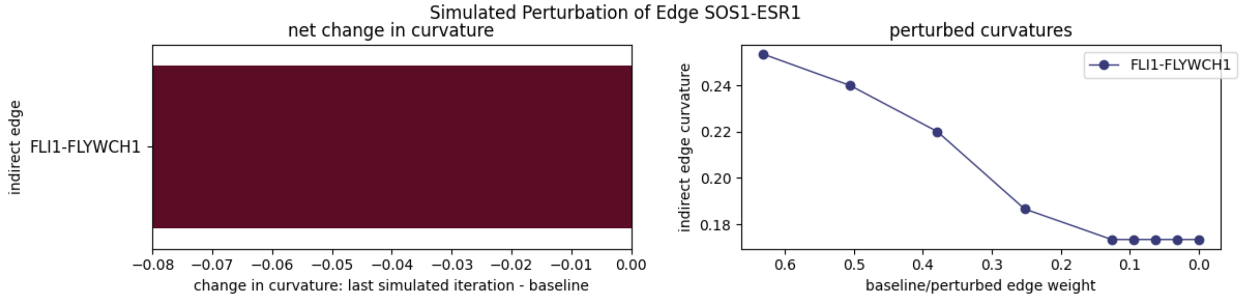


Figure S23: Perturbing edge *SOS1-ESR1*.

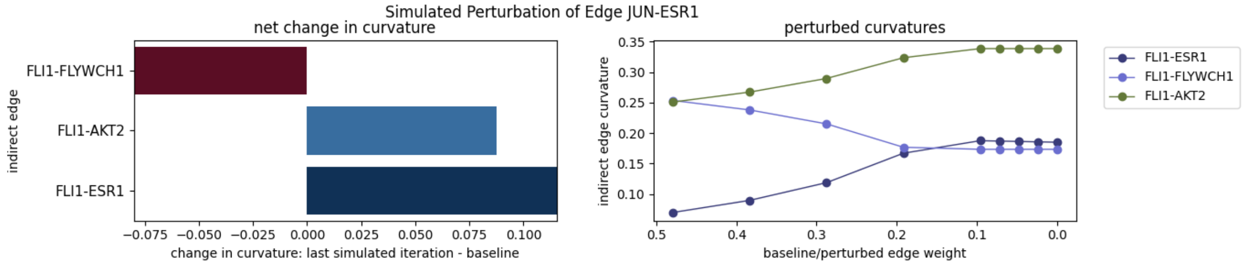


Figure S24: Perturbing edge *JUN-ESR1*.

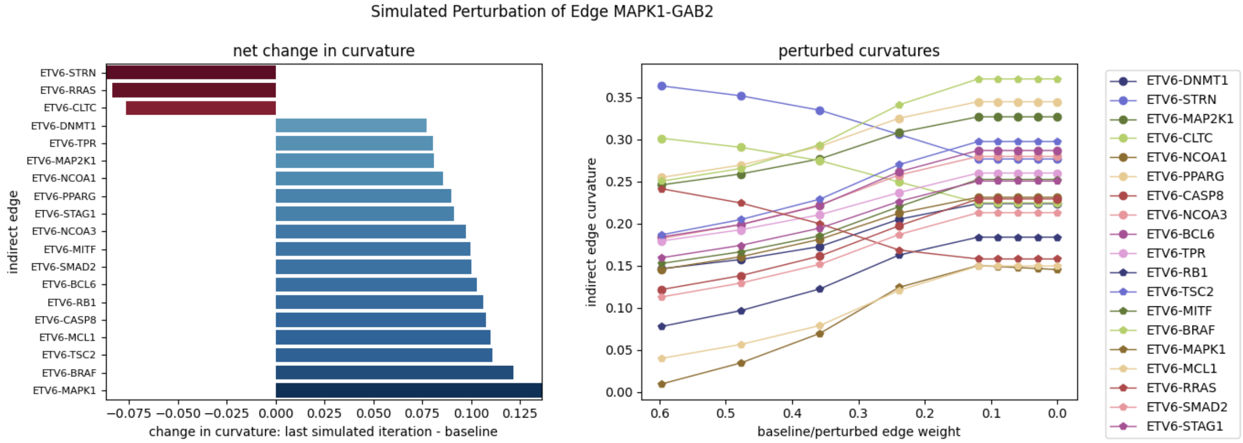


Figure S25: Perturbing edge *MAPK1-GAB2*.

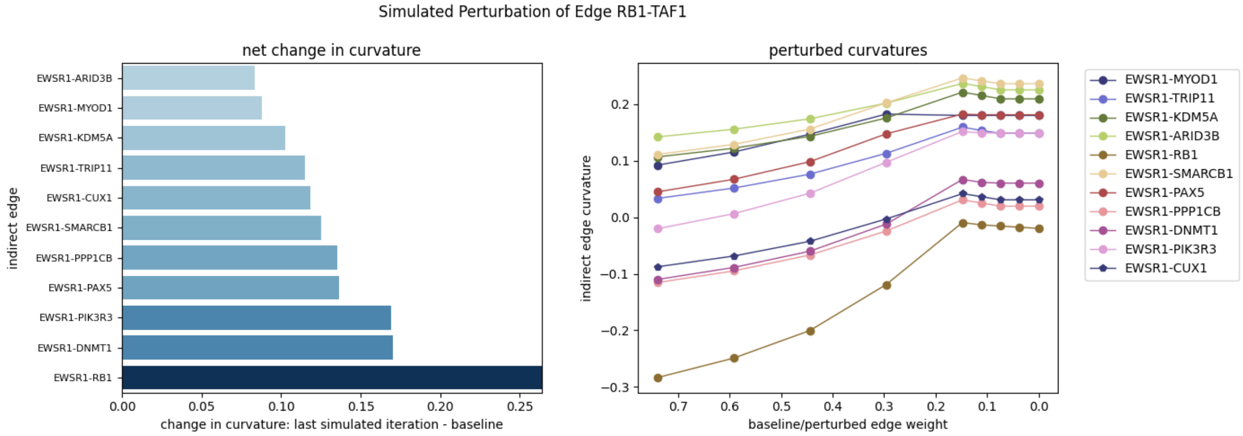


Figure S26: Perturbing edge *RB1-TAF1*.

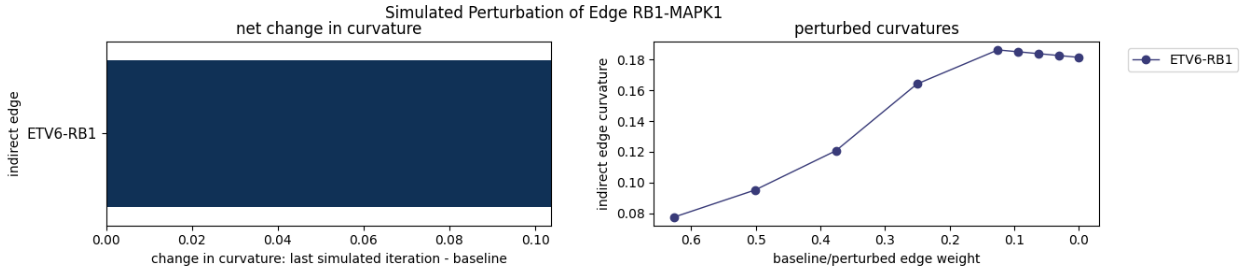


Figure S27: Perturbing edge *RB1-MAPK1*.

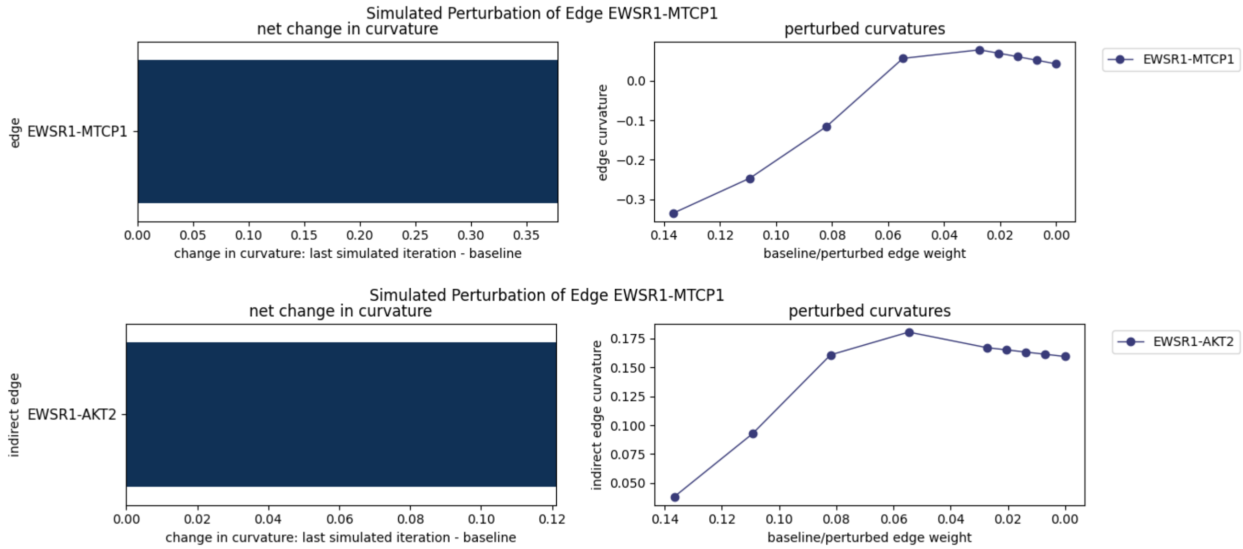


Figure S28: Perturbing edge *EWSR1-MTCP1*. (Top) Direct affected interactions, i.e., edges. (Bottom) Indirect affected interactions.

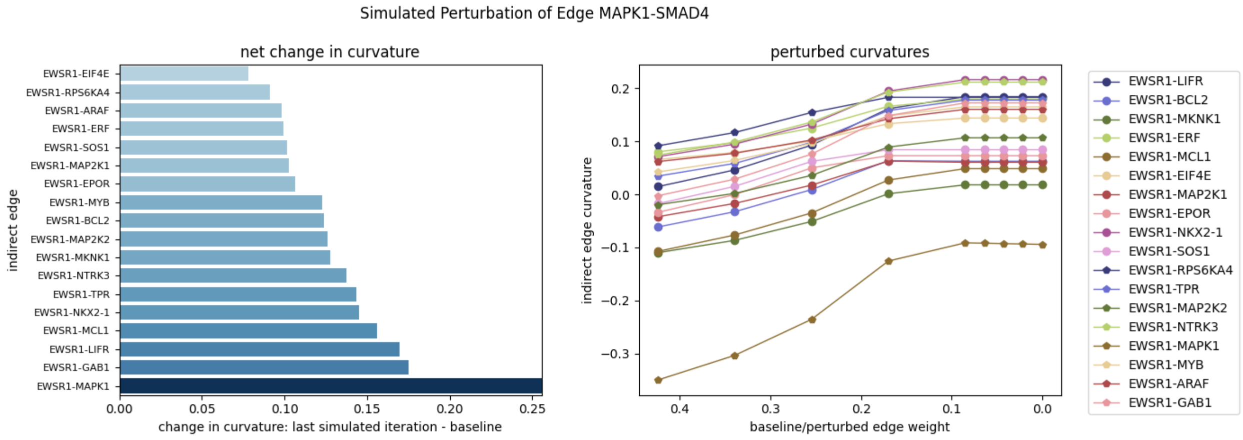


Figure S29: Perturbing edge *MAPK1-SMAD4*.

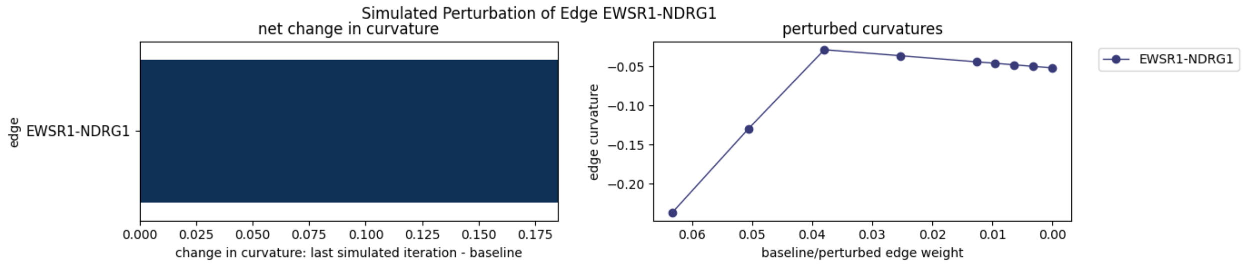


Figure S30: Direct interactions affected by perturbing edge *EWSR1-NDRG1*.

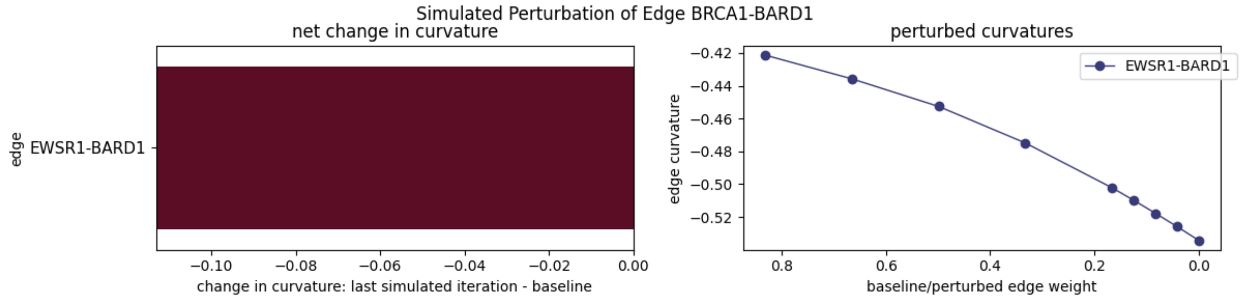


Figure S31: Direct interactions affected by perturbing edge *BRCA1-BARD1*.

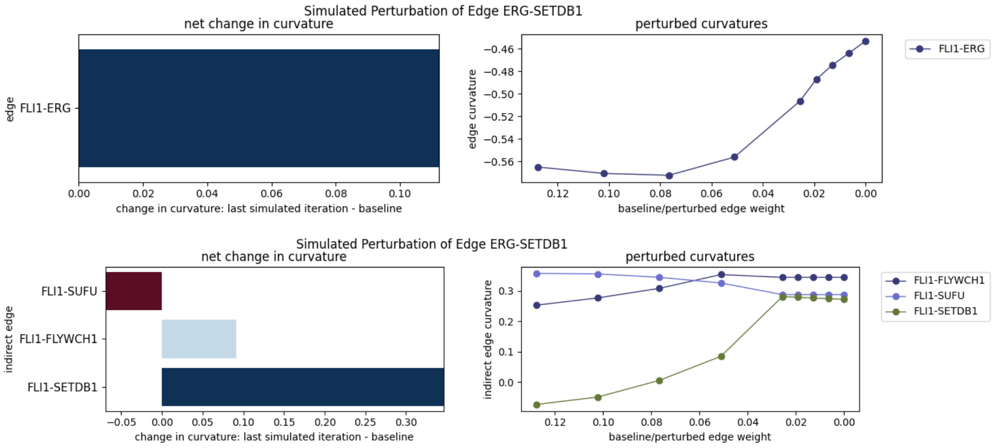


Figure S32: Perturbing edge *ERG-SETBP1*. (Top) Direct affected interactions, i.e., edges. (Bottom) Indirect affected interactions.

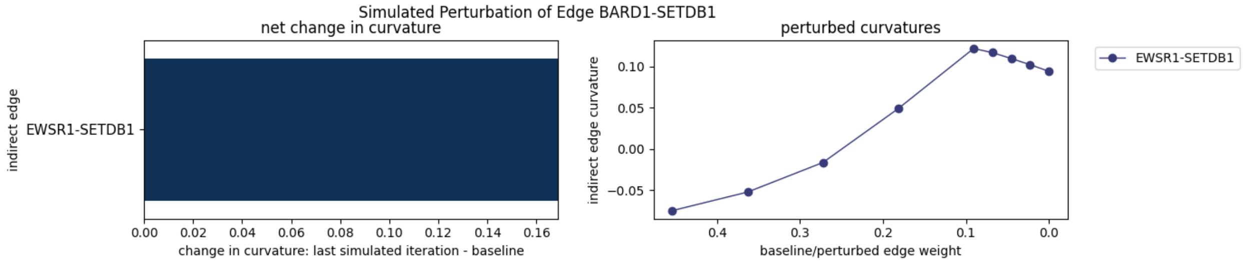


Figure S33: Direct interactions affected by perturbing edge *BARD1-SETDB1*.

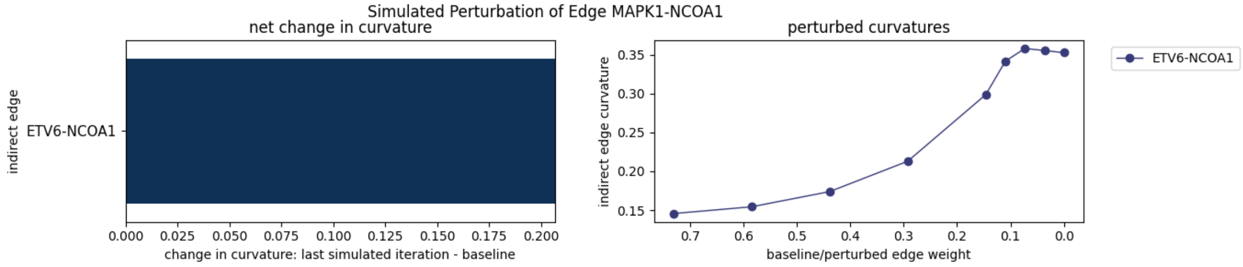


Figure S34: Direct interactions affected by perturbing edge *MAPK1-NCOA1*.
